# HCN channels enhance robustness of patterned activity propagation in heterogeneous conductance-based ring networks

**DOI:** 10.1101/2023.11.28.568983

**Authors:** Divyansh Mittal, Rishikesh Narayanan

**Affiliations:** Cellular Neurophysiology Laboratory, Molecular Biophysics Unit, Indian Institute of Science, Bangalore, India

**Keywords:** biophysical network models, continuous attractor network, degeneracy, heterogeneities, negative feedback, neuronal resonance, robustness, stability

## Abstract

Continuous attractor network (CAN) models lend a powerful framework that has provided deep insights about several aspects of brain physiology. However, most CAN models employ homogeneous, rate-based or artificially spiking neurons with precisely structured synaptic connectivity, precluding detailed analyses of the impact of specific neural-circuit components and associated heterogeneities on CAN dynamics. To address this caveat, we built populations of tunable and scalable conductance-based, physiologically constrained, ring network models consisting of distinct rings of excitatory and inhibitory neurons. We assessed the network for its ability to sustain robust propagation of patterned activity across the rings. First, in homogeneous ring networks, we found that robust activity propagation could be sustained through several different combinations of synaptic weights, demonstrating synaptic degeneracy in the emergence of robust activity propagation. We incorporated intrinsic heterogeneity through randomized perturbations to ion channel parameters of all neurons and synaptic heterogeneity by adding jitter to the Mexican-hat connectivity between inhibitory neurons. We found the number of networks exhibiting robust propagation of patterned activity to reduce with increase in the degree of synaptic or intrinsic heterogeneities. Motivated by the ability of intrinsic neuronal resonance to stabilize heterogeneous rate-based CAN models, we hypothesized that increasing HCN-channel (a resonating conductance) density would stabilize activity propagation in heterogeneous ring networks. Strikingly, we observed that increases in HCN-channel density resulted in a pronounced increase in the proportion of heterogeneous networks that exhibited robust activity propagation, across multiple trials and across three degrees of either form of heterogeneity. Together, heterogeneous networks made of neurons with disparate intrinsic properties and variable HCN channel densities yielded robust activity propagation, demonstrating intrinsic degeneracy in the emergence of robust activity propagation. Finally, as HCN channels also contribute to changes in excitability, we performed excitability-matched controls with fast HCN channels that do not introduce resonance. We found that fast HCN channels did not stabilize heterogeneous network dynamics over a wide range of conductance values, suggesting that the slow negative feedback loop introduced by HCN channels is a critical requirement for network stabilization. Together, our results unveil a cascade of degeneracy in ring-network physiology, spanning the molecular-cellular-network scales. These results also demonstrate a critical role for the widely expressed HCN channels in enhancing the robustness of heterogeneous neural circuits by implementing a slow negative feedback loop at the cellular scale.

## INTRODUCTION

The brain is a complex system that creates order in its representations of the world and governs intricate functions through collective interactions among simpler elements. These simpler elements manifest functional segregation but show increasing degrees of functional integration when collective interactions among them drive complex functions (Edelman and Gally, 2001). The continuous attractor network (CAN) is an outstanding example of a system that manifests such emergence of complex function through intricate interactions among simpler elements (Khona and Fiete, 2022). Each neuronal element that is part of CANs is a functionally segregated unit, but they manifest functional integration through synaptic interactions towards the emergence of continuous attractor dynamics. The mechanistic details of how neuronal interactions yield continuous attractor dynamics, the specific connectivity constraints that are essential for CAN emergence, and the several outstanding utilities of CAN networks in behavioral contexts are well studied (Redish et al., 1996; Samsonovich and McNaughton, 1997; Seung et al., 2000; Renart et al., 2003; Song and Wang, 2005; Wills et al., 2005; Burak and Fiete, 2009; Knierim and Zhang, 2012; Couey et al., 2013; Domnisoru et al., 2013; Schmidt-Hieber and Hausser, 2013; Yoon et al., 2013; Widloski and Fiete, 2014; Seelig and Jayaraman, 2015; Knierim and Neunuebel, 2016; Shay et al., 2016; Kim et al., 2017b; Kim et al., 2019; Stella et al., 2020; Spalla et al., 2021; Tukker et al., 2021; Khona and Fiete, 2022).

Despite this, owing predominantly to computational complexity, there are two major caveats in existing analyses of CAN models. First, many CAN models assume idealized scenarios where the simpler elements (the neurons) are repeating homogeneous replicates with precisely defined spatially replicating interactions (synaptic connections) between them, manifesting no heterogeneities. However, heterogeneities are ubiquitous across scales of analysis in biological systems (Renart et al., 2003; Gjorgjieva et al., 2016; Cembrowski and Spruston, 2019; Mishra and Narayanan, 2019; Rathour and Narayanan, 2019; Mittal and Narayanan, 2021; Perez-Nieves et al., 2021; Mittal and Narayanan, 2022; Seenivasan and Narayanan, 2022; Shridhar et al., 2022; Segal et al., 2023). Hence, one of the fundamental requirements for extending the CAN framework to biological substrates is to assess the impact of neural-circuit heterogeneities on CAN model performance. If the continuous attractor dynamics were entirely reliant on artificial homogeneity of neuronal and synaptic elements, then the extendibility of the CAN framework to inherently heterogeneous biological systems becomes impossible.

Second, most CAN models do not consider the complexities and the heterogeneities associated with the *simpler elements* that form this network. Individual neurons used in such models often exhibit rate-based dynamics or are artificially spiking neurons that lack details of the specific intrinsic electrophysiological properties exhibited by their biological counterparts. Although the different neurons that make the network are simple elements from the network perspective, they are themselves complex systems that are built from heterogeneous elements. Specifically, characteristic electrical properties of neurons are governed by a variety of molecules (ion channels, pumps, and buffers) that are subtype specific. Importantly, these characteristic physiological properties can be achieved through several non-unique combinations of these constitutive components, a phenomenon that has been referred to as degeneracy (Edelman and Gally, 2001; Rathour and Narayanan, 2019; Goaillard and Marder, 2021). Thus, neural heterogeneities do not just span single-neuron physiological properties, but also in the molecular components that yield these signature characteristics. If there are several non-unique routes to constructing neurons with specific characteristic physiological properties, the impact of such molecular heterogeneities on robust network function becomes an important question to consider. Additionally, as physiological plasticity, behaviorally driven neuromodulation, or pathological conditions affect specific *molecular* components, there is a need to account for subtype-specific differences in molecular composition and associated heterogeneities in assessing continuous attractor networks. If the molecular components of different neuronal subtypes or the neuron-to-neuron variability in these components are not accounted for, experiments or models would not accurately assess the impact of physiological or pathological perturbations on CAN output or associated behavior.

In rectifying these two prominent caveats associated with several CAN models, we built conductance-based excitatory (*E*)–inhibitory (*I*) ring networks endowed with cellular and molecular heterogeneities and assessed robust propagation of patterned activity across the ring network. The fundamental question we address here is on how complex networks of neurons robustly execute their collective function in the presence of heterogeneities in the cells that they are composed of as well as in the molecules that yielded those cells. Addressing this question effectively requires heterogeneous populations of neurons of each neuronal subtype (*E* and *I*), all of which manifest respective characteristic physiological properties but through different cell-type specific molecular combinations. Thus, we first constructed heterogeneous populations of excitatory (*n*=449 neurons from (Mittal and Narayanan, 2018)) and inhibitory (*n*=930 neurons, constructed here) neurons, each independently manifesting parametric degeneracy on cellular-scale physiology. We built homogenous as well as heterogeneous ring networks using neurons from these populations and assessed their functional robustness to different kinds of perturbations. Our ring network is a one-dimensional adaptation of the 2D network proposed by Burak and Fiete (Burak and Fiete, 2009), built with heterogeneous conductance-based models for the distinct *E* and *I* neuronal populations.

We found that introduction of heterogeneities in synaptic connectivity or intrinsic properties into the network severely hampered the ability of the ring network to sustain patterned activity propagation. Motivated by the ability of intrinsic neuronal resonance to stabilize rate-based CAN networks (Mittal and Narayanan, 2021), we asked if increasing resonance frequency by increasing HCN channels could enhance robustness in heterogeneous ring networks. Strikingly, increases in HCN-channel density, in inhibitory but not excitatory neurons, increased the proportion of heterogeneous networks that exhibited robust activity propagation. Finally, as HCN channels also contribute to changes in excitability, we performed excitability-matched controls with fast HCN channels that do not introduce resonance. We found that fast HCN channels did not stabilize CAN model dynamics over a wide range of conductance values, suggesting resonance and the associated slow negative feedback loop (that suppresses low frequency activity) as critical requirements for stabilization of ring network dynamics. Together, our results point to a novel role for the widely expressed resonating conductances in enhancing functional robustness of heterogeneous neural circuits. Our analyses also clearly demonstrate how ion channels and intrinsic properties of constituent neurons can guide and stabilize network-level computation through several non-unique routes, pointing to manifestation of degeneracy spanning multiple scales of neural function.

## RESULTS

### Heterogeneous conductance-based model populations of physiologically validated excitatory and inhibitory neurons

Our first step in building a conductance-based Excitatory (*E*)–Inhibitory (*I*) ring network was to construct biophysically and physiologically relevant models of excitatory and inhibitory neurons that would together form the ring network. Neuronal intrinsic properties are mediated by passive neuronal properties and active components such as voltage- and calcium-gated ion channels. Thus, our goal was to build conductance-based models of *E* and *I* neuronal populations using characteristic ion channels expressed in these neuronal subtypes and validated against their respective characteristic physiological properties. We used a pre-validated heterogeneous model population of medial entorhinal cortex layer II stellate cells (*n* = 449), with 9 active conductances, as *E* neurons of the conductance-based ring network. This model population was built using a multi-parametric multi-objective stochastic search (MPMOSS) algorithm and was validated against characteristic electrophysiological properties of entorhinal cortical stellate cells (Mittal and Narayanan, 2018).

We used an independent MPMOSS algorithm to generate a population of inhibitory cortical neurons (fast-spiking basket cells). Towards this, we hand-tuned a conductance-based base model endowed with active and passive properties of cortical interneurons (Fig. 1*A–G*). The model was constructed with passive leak channels and 5 different active ion-channel subtypes derived from cortical interneurons. The base model parameters were tuned to match 10 different characteristic electrophysiological measurements from cortical interneurons (Tzilivaki et al., 2019). The characteristic properties of cortical interneurons that were matched included the prominent fast afterhyperpolarization (*AHP*; Fig. 1*G*), the high firing rate (*f*_250_, the firing rate response to a pulse current of 250 pA; Fig. 1*E*), and narrow action potential half width (*T*_$%&’_; Fig. 1*G*). The 6 active and passive channel parameters (maximal conductance of the leak channels, specified as specific membrane resistance, and 5 active channels) that were tuned to obtain required model outcomes are mentioned in Table 1.

**Figure 1:**
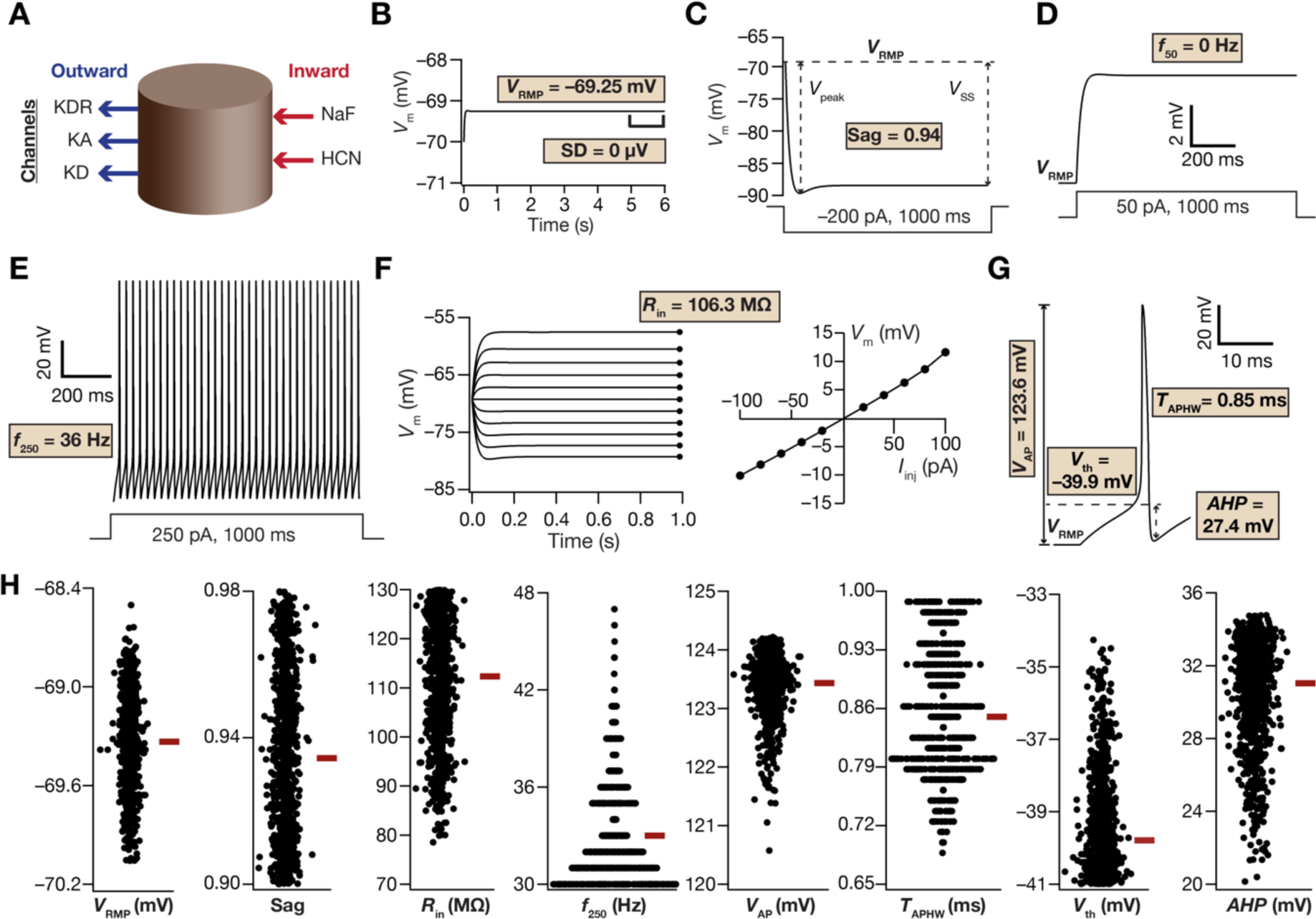
Cortical interneuron base model and the distribution of electrophysiological measurements from valid cortical interneuron model population obtained using MPMOSS. (A) Schematic representation of a single compartment model for cortical interneuron specifying inward (inward arrows) and outward (outward arrows) currents. (B*–*I) The 10 physiologically relevant measurements (highlighted within boxes showing the symbol along with the measurement in the base model) used to characterize cortical interneurons. (B) Resting membrane potential (*V*RMP) and its standard deviation (SD) were computed by taking the mean and standard deviation, respectively, of the membrane potential between 5–6 s duration (window specified in the figure) when no current was injected. All the other measurements were performed after the model settled at its *V*RMP at 6 s. (C) Sag ratio (Sag) was measured as the ratio of the steady-state membrane potential deflection (*V*SS) to peak membrane potential deflection (*V*peak) in the voltage response of the model to a hyperpolarizing step current of 200 pA for a duration of 1000 ms. (D*–*E) Voltage response of the model to a step current of 50 pA (D) or 250 pA (E) for a stimulus duration of 1000 ms was used to measure the number of action potentials (*f*50 or *f*250) elicited for the respective current injection. (F) Input resistance (*R*in) computation. *Left*, 1000 ms long step currents from –100 pA to 100 pA were injected into the cell in steps of 20 pA to record the steady-state voltage response (black circles at the end of each trace). *Right*, Steady-state voltage response *vs.* injected current (*V–I*) plot obtained from the traces on the left panel. The slope of a linear fit to the *V–I* plot defined *R*in. (G) Amplitude of action potential (*V*AP) was measured as the difference between the peak voltage achieved during an action potential and *V*RMP. Threshold of action potential (*V*th) was the membrane potential at which the first derivative of action potential crosses 20 V/s. Action potential width at half maximum (*T*APHW) defined the width of action potential at half point between maximum amplitude achieved by action potential and action potential threshold (*V*th). Afterhyperpolarization (*AHP*) is the difference in membrane potential between action potential threshold (*V*th) and minimum membrane potential achieved by the end of action potential. (H) Heterogenous distribution of physiologically relevant measurements in the valid cortical interneuron models obtained after a multi-parametric multi-objective stochastic search. Bee-swarm plots depicting the distribution of 8 measurements in the 930 valid models. The red rectangle adjacent to each plot represents the respective median value. The standard deviation (SD) measurement depicting stability of RMP was uniformly low and *f*50 was identically zero across all models, and therefore are not depicted as plots here.

**Table 1.** Range of parameters used in generating the population of cortical interneuron models.

| No. | Parameter | Description | Min | Max |
| --- | --- | --- | --- | --- |
| 1 | $g^{\text{NaF}}$ (mS/cm<sup>2</sup>) | Maximal conductance of fast sodium (NaF) channel | 30 | 150 |
| 2 | $g^{\text{KDR}}$ (mS/cm<sup>2</sup>) | Maximal conductance of delayed rectifier potassium (KDR) channel | 3 | 30 |
| 3 | $g^{\text{HCN}}$ (μS/cm<sup>2</sup>) | Maximal conductance of Hyperpolarization-activated cyclic-nucleotide-gated (HCN) channel | 1 | 10 |
| 4 | $g^{\text{KA}}$ (mS/cm<sup>2</sup>) | Maximal conductance of A-type potassium (KA) channel | 0.25 | 2.5 |
| 5 | $g^{\text{KD}}$ (μS/cm<sup>2</sup>) | Maximal conductance of D-type potassium (KD) channel | 5 | 50 |
| 6 | $R_m$ (kΩ cm<sup>2</sup>) | Specific membrane resistance | 15 | 40 |

An unbiased stochastic search, looking for other valid cortical interneuronal models, was performed around the parametric values associated with the base model. Specifically, a unique random value was independently selected for each of the 6 parameters from respective uniform distributions that spanned around their respective base values (Table 1). This set of six random values acted as the substrate for a model that was constructed. We generated 10,000 such randomized models and validated the 10 different measurements obtained from them with their electrophysiological counterparts (Table 2). 930 of the 10,000 models that satisfied all ten electrophysiological bounds were declared as valid models of cortical interneurons. Importantly, the resultant model population exhibited wide-spread heterogeneity in their electrophysiological measurements (Fig. 1*H*), within their respective characteristic ranges (Table 2).

**Table 2.** Physiologically relevant range of cortical interneuron measurements. Experimental bounds on each intrinsic measurement involved in the validation process of stochastically generated models. While the constraint on the standard deviation of resting membrane potential ensures that there are no membrane potential oscillations at rest, the rest of the bounds were derived from previous electrophysiological measurements (Tzilivaki et al., 2019).

| No. | Measurements | Lower Bound | Upper Bound |
| --- | --- | --- | --- |
| 1 | Resting membrane potential, $V_{\text{RMP}}$ (mV) | -75 | -65 |
| 2 | Standard deviation of membrane potential (Resting), SD (mV) | 0 | 1 |
| 3 | Sag ratio, $Sag$ | 0.90 | 0.98 |
| 4 | Input resistance, $R_{\text{in}}$ (MΩ) | 70 | 130 |
| 5 | Action potential threshold, $V_{\text{th}}$ (mV) | -33 | -41 |
| 6 | AP amplitude, $V_{\text{AP}}$ (mV) | 80 | N/A |
| 7 | Action potential width at half maximum, $T_{\text{APHW}}$ (ms) | N/A | 1 |
| 8 | Afterhyperpolarization, $AHP$ (mV) | 20 | N/A |
| 9 | Number of APs for a 50 pA step current for 1000 ms, $f_{50}$ (Hz) | 0 | 0 |
| 10 | Number of APs for a 250 pA step current for 1000 ms, $f_{250}$ (Hz) | 30 | 50 |

We next asked if there were strong correlations between the electrophysiological measurements that defined these cortical interneuron models. Strong correlations between these measurements would suggest the presence of a smaller set of core measurements, with the other measurements relegated to being redundant in terms of placing strong constraints on the solution space. Strong correlations between measurements also would suggest the dependence of these measurements on a smaller set of ion-channels. To find relationships across measurements, we computed pairwise Pearson’s correlation coefficients across all the measurements for the valid model population of cortical interneurons. We found that while most of the measurements did not exhibit strong correlations with each other, a few measurements which are known to be critically controlled by similar set of ion-channels depicted strong pair-wise correlations (Fig. 1–figure supplement 1). For example, measurements like *Sag* and *V*RMP, which are known to be critically reliant on HCN channels (Mishra and Narayanan, 2023) showed strong pair-wise correlations. These observations suggest that the different measurements characterized different aspects of interneuron physiology and there was a lack of strong dependencies in most measurements.

Together, an independent MPMOSS algorithm that was biophysically and physiologically constrained by cortical interneuron characteristics yielded a heterogeneous population of model neurons, which formed the *I* neurons of our conductance-based ring network.

### Ion-channel degeneracy in the model population of cortical interneurons

Were the parameters underlying this population of models that manifested characteristic cortical interneuron properties clustered in the parametric space? Were there strong constraints on these parameters to yield a physiologically valid model of cortical interneurons? Did these neurons exhibit ion-channel degeneracy, whereby disparate combinations of underlying parameters yield cortical interneurons with similar physiological characteristics? To address these questions, we first picked 5 models, out of the 930 valid ones, that were endowed with very similar measurements (Fig. 2*A–F*). We asked if the parameters associated with these models were similar (clustered) or manifested heterogeneities. We found that the parameters associated with these functionally similar models were distributed across respective minimum and maximum for each parameter (Fig. 2*G*). These observations provided an illustrative line of evidence for the expression of degeneracy in cortical interneurons, towards the manifestation of characteristic physiological properties. We assessed the parametric ranges of all valid models and found that their underlying parameters covered their respective ranges (Fig. 2*H*, bottommost histogram panel). Together, our analyses demonstrate that various combinations of parametric values, spanning wide ranges, could yield cortical interneuron models with characteristic physiological properties.

**Figure 2:**
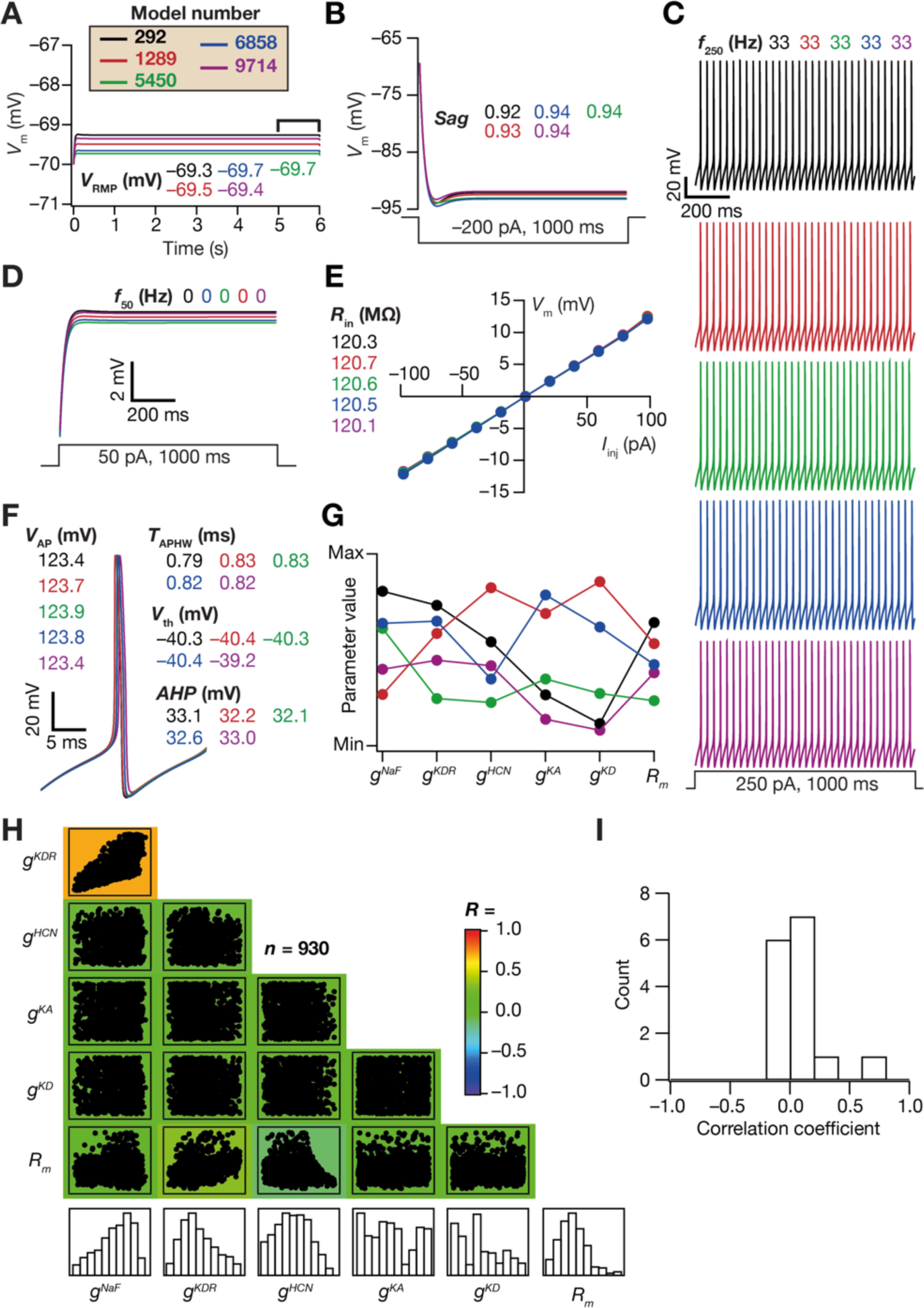
Disparate combinations of model parameters resulted in similar physiological measurements in 5 valid cortical interneuron models pointing to the expression of cellular-scale degeneracy. (A–F) Voltage traces and 10 physiologically relevant measurement values from 5 randomly chosen valid models obtained after MPMOSS. The measurements reported are resting membrane potential (*V*RMP) and its standard deviation (SD) which is zero for all the models (A), Sag ratio (B), Number of action potentials for a step current injection of 250 pA for 1000 ms (*f*250) (C), Number of action potentials for a step current injection of 50 pA for 1000 ms (*f*50) (D), Input resistance (*R*in) (E), amplitude of action potential (*V*AP), action potential threshold (*V*th), action potential width at half maximum (*T*APHW), and afterhyperpolarization (*AHP*) (F). (G) Normalized values of each of the 6 parameters that were employed in the generation of these models, shown for the 5 randomly chosen models depicted in A*–*F. Each parameter was normalized by the respective minimum and maximum values that bound the stochastic search for that parameter (Table 1). Parameters associated with corresponding model traces in panels A*–*F are depicted with identically color-coded markers in panel G. (H) Pair-wise scatter plots between 6 parameters for all the valid cortical interneuron models (*n*=930). The bottom most row depicts the histograms for the corresponding parameters in the valid heterogeneous population. Individual scatter plots are overlaid on a heat map that depicts the pair-wise correlation coefficient computed for that scatter plot. (I) Distribution of correlation coefficient values of 15 unique pairs between the 6 parameters from scatter plots in (H).

Were there strong interdependencies across parametric combinations that yielded the valid model population? To assess this, we computed pair-wise Pearson’s correlation coefficient (*R*) values across all parametric values of the valid models (Fig. 2*H*). We found weak pair-wise correlations between most parameters (Fig. 2*H–I*) except for a relatively strong correlation (*R*=0.7) between maximal conductance values of the two spike-generating conductances: delayed rectifier potassium (*g*^()*^) and fast sodium (*g*^+,-^). Together, these observations demonstrated the manifestation of ion-channel degeneracy in the interneuronal model population that was endowed with widespread distributions of and weak pairwise correlations across underlying parameters.

### Stable and robust propagation of patterned activity across the conductance-based E–I ring network

These previous steps provided us independent heterogeneous populations of cortical excitatory (*E*) and inhibitory (*I*) neurons. Towards realizing our principal goal of understanding the impact of neural-circuit heterogeneities on patterned activity propagation in *E–I* ring networks, we first built a homogeneous *E–I* ring network where heterogeneities would be introduced progressively. The network was made of two rings: an *E*-ring made of 80 *E* neurons and an *I*-ring constructed with 20 *I* neurons (Fig. 3*A*). In building a homogeneous ring network model, we first chose one model neuron each from both excitatory and inhibitory valid model populations, with medium excitability (firing rate of chosen *E* neuron to a 250 pA current injection=11 Hz; firing rate of chosen *I* neuron to a 250 pA current injection=37 Hz). We replicated the chosen *E* neuron 80 times to construct the homogeneous *E* ring. The chosen *I* neuron was replicated 20 times to build the homogeneous *I* ring. *I* neurons made inhibitory GABAergic synapses with each other as well as with *E* neurons in a manner that followed distance-dependent Mexican-hat connectivity profiles. The connectivity between *E* neurons to *I* neurons was all-to-all and glutamatergic in nature, with no connectivity among *E* neurons (Fig. 3*A*). All excitatory connections were implemented using AMPA receptor and inhibitory connections were realized using GABAA receptors. In this homogeneous ring network, we hand-tuned the network parameters associated with the coupling of afferent inputs, the strengths and shapes of Mexican-hat connectivity, and the strength of all-to-all *E*-to-*I* connectivity towards achieving patterned activity propagation across the network.

**Figure 3:**
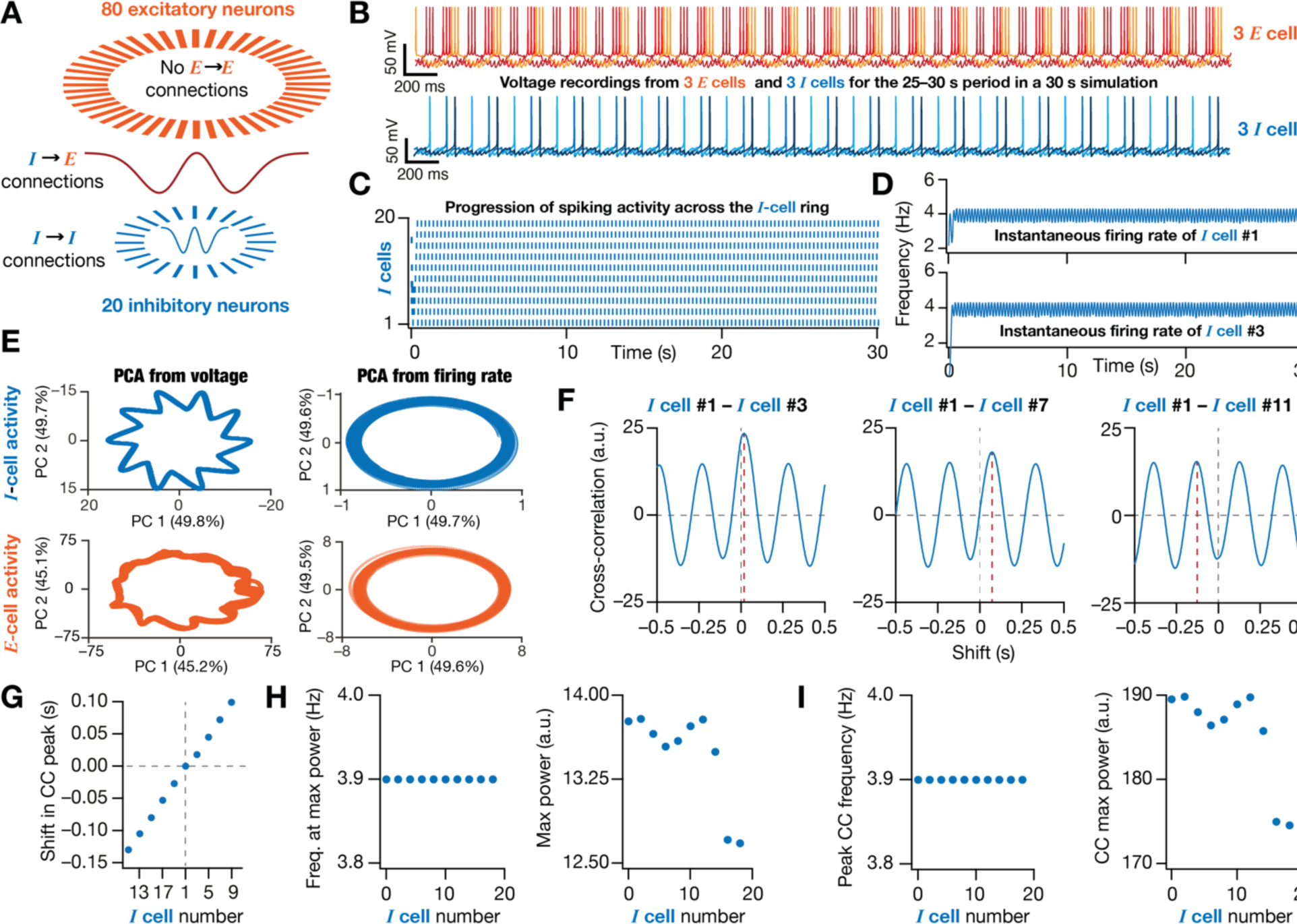
Conductance-based ring network model showing patterned activity propagation. (A) Graphical representation of network architecture with excitatory (*E* cell) and inhibitory (*I* cell) neurons in the ratio of 4:1 (default, 80 excitatory and 20 inhibitory conductance-based model neurons). The synaptic connectivity among inhibitory neurons follows distance-dependent periodic Mexican-hat connectivity. Excitatory neurons are not connected with each other. The inhibitory to excitatory neurons connections also follow distance-dependent periodic Mexican-hat connectivity. Connections between excitatory neurons to inhibitory neurons were all-to-all. (B) Voltage traces from 3 excitatory and 3 inhibitory neurons in the network showing pattern propagation of activity across the rings of *E* (top) and *I* (bottom) cells. The temporal shifts in the time of spiking of different *E* as well as *I* cells may be noticed. (C) Raster plot showing activity of 10 inhibitory neurons (total *n* = 20; shown are every alternate neuron) exhibiting patterned activity propagation across the ring. Each tick represents a spike in the neuron specified by the column. (D) Example instantaneous firing rate of two *I* cells, computed by convolving the binarized temporal spike trains (in panel C) of each cell with a Gaussian kernel of standard deviation, *σ*_*FR*_=100 ms. (E) Population dynamics of *I*-cells (top) and *E*-cells (bottom) manifested low-dimensional ring manifolds. Principal component analysis (PCA) was performed either on the smoothed (Gaussian *σ*=40 ms) voltage traces (left) or instantaneous firing rate (right; examples in panel D) spanning all *I* or *E* cells in the network. The variance explained by the first two principal components (PC) were large enough for all cases (the variance explained by each PC is given within parentheses). (F) Example cross-correlogram (CC) between different *I* cells in the network with reference to *I* Cell #1. The progressive shift in the peak of cross-correlogram (shown with red dotted line) indicates propagating activity across the network. (G) Plot showing the time points at which the cross-correlograms (with reference to *I* Cell #1, from panel F) peaked for each *I* cell. The cells are arranged to depict the patterned propagation of the activity bump across the ring network. This progressive shift in the temporal location of the peak of the cross-correlogram depicts attractor movement over the inhibitory neural lattice. (H) Average firing frequency (*Left*) and maximum power (*Right*) from spectral analysis of the activity of each neuron in the network (examples in panel D). (I) Frequency of patterned activity (*Left*), and maximum power (*Right*) computed from spectral analysis of cross-correlograms of *I* Cell #1 with all the other cells (examples in panel F).

In this hand-tuned *E–I* ring network, based on neuronal voltage responses and raster plots spanning different cells, we observed stable propagation of patterned activity across the *I*-ring (Fig. 3*B–D*) as well as the *E*-ring (Fig. 3–figure supplement 1*B–C*). Given the connectivity patterns in our *E–I* ring network (Fig. 3*A*) where *E*-to-*E* connections are absent, patterned activity in the network emerges as a consequence of the *I*-to-*I* and *I*-to-*E* Mexican-hat connectivity patterns, thus making the *I* ring to be central for patterned activity propagation across both rings. Importantly, when we visualized population activity across the *I*-cell or the *E*-cell networks in a reduced dimensional space, we found the manifestation of a low-dimensional ring manifold in the population activity dynamics (Fig. 3*E*). This was irrespective of whether we used smoothed voltage traces or instantaneous firing rates as representatives of population neural activity (Fig. 3*E*). These observations provide an important line of evidence for the manifestation of a ring attractor in our *E–I* ring network.

To quantify such patterned propagation, we developed a set of metrics that exploited two specific characteristic features of patterned activity propagation across the ring. First, with robust propagation of patterned activity, the neuronal firing would be repetitive, implying that the activity bump returns to the same neuron in periodic manner owing to ring architecture. We quantified this aspect of patterned activity by assessing periodicity in the Fourier magnitude spectrum of the instantaneous activity (Fig. 3*D*) as well as the cross-correlogram (CC) of instantaneous activity across different cells (Fig. 3*F*). Second, activity propagation would be temporally sequential along the ring, implying that the shift in the temporal position of the activity bump between two neurons in a ring would depend on the distance between the two neurons in the ring. We quantified this by computing the cross-correlogram of instantaneous firing rate of each cell in the ring with respect to an arbitrarily chosen reference cell (Fig. 3*F*). We computed the shift in the peak location of each of these correlograms and plotted them as the function of sequential cell position in the ring (Fig. 3*G*). Robust propagation of patterned activity along the network would manifest a progressive shift in the position of the peak CC value, providing a non-zero slope for the plot between the CC peak location and cell position (Fig. 3*G*).

We computed five distinct metrics to assess stability of patterned activity propagation from the instantaneous firing rate as well as cross-correlograms associated with all cells in the ring. As mentioned above, a non-zero slope of the plot between the shift in CC peak and cell position indicated robust propagation of patterned activity across the ring (Fig. 3*G*; CC slope=0.0252). We computed the maximum power (*P*_./0_) as well as the frequency at which this peak occurred (*f*_12/3_) from the Fourier magnitude spectrum of the instantaneous firing rate of each cell (Fig. 3*H*). The value *f*_12/3_ represented the frequency at which activity returned to the same cell in the ring, while *P*_./0_ indicated the strength of the patterned activity (Fig. 3*H*). In a ring network sustaining robust propagation of patterned activity, there would be minimal variability in *P*_./0_ and *f*_12/3_across different cells in the network (Fig. 3*H*). Thus, we computed the coefficient of variation (CV) of *P*_./0_ and *f*_12/3_ to assess similarity of patterned activity across all cells in the ring. With a similar rationale, we calculated the maximum power (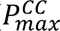 and the frequency of peak power (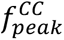 for the cross-correlograms associated with each cell in the ring (with reference to a reference cell). We observed that the CV of these quantities were low when the ring sustained robust propagation of patterned activity (Fig. 3*H–I*; CV of *P*_./0_= 0.0324; CV of *f*_12/3_=0 Hz; CV of 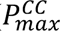 = 0.0314; CV of 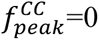 =0 Hz). These five metrics derived from the instantaneous firing rates and the cross-correlograms across cells quantified stable propagation of patterned activity across the *I* ring (Fig. 3) as well as the *E* ring of the network (Fig. 3–figure supplement 1).

Together, we built a homogeneous conductance-based *E–I* ring network, with repeating *E* and *I* neurons with identical connectivity patterns, that sustained robust propagation of patterned activity along both rings. We also defined different metrics which could be together used for quantifying patterned activity propagation in these conductance-based networks.

### Robustness of patterned propagation of activity across the ring to changes in different network parameters

How robust was patterned activity propagation to changes in the size of the network? We progressively increased the size of our homogeneous *E–I* ring network, while maintaining the connectivity patterns and *I:E* neuronal ratio, to assess the ability of the network to sustain patterned activity propagation across the ring (Fig. 3–figure supplement 2). We found our conductance-based ring network to be scalable when the network size was increased, robustly sustaining propagation of patterned activity across the ring (Fig. 3–figure supplement 2). However, the minimum size at which patterned propagation of activity could be observed was in the default 100 neuron ring network, with a network with lower size incapable of sustaining patterned activity propagation (Figure 3–figure supplement 2). These observations are consistent with a threshold on the minimum network size required for sustaining continuous attractor dynamics and activity bump propagation (Burak and Fiete, 2009; Mittal and Narayanan, 2021).

In constructing the homogeneous ring network, we had picked one each of the valid *E* and *I* neurons, from their respective populations, with medium excitability (firing rate) and used them for constructing the *E* and *I* rings, respectively. How did the intrinsic excitability of the individual neurons affect the ability of a *homogeneous* ring network to sustain patterned activity propagation? To assess this, we constructed homogeneous ring networks with different *E* and *I* neurons. These neurons were picked from their respective valid model populations, but had either low or high excitability compared to the neurons we picked earlier. The chosen low excitability *E* and *I* neurons had their firing rate for 250 pA current injection, *f*_250_, as 7 and 31 Hz, respectively. Their high excitability counterparts had *f*_250_as 15 and 46 Hz, respectively. In building a homogeneous ring network with low excitability *E* neurons, we replaced all the medium-excitability *E* neurons in the network identically by the chosen low-excitability *E* neurons. Thus, the network continued to be homogeneous with all neurons in a ring identical, showing low (L), medium (M), or high (H) excitability. This design yielded nine different combinations corresponding to three excitability levels each for *E* and *I* neurons (Fig. 4). The network shown in Fig. 3 corresponds to the Medium-excitability *I*–Medium-excitability *E* (MI–ME) configuration.

**Figure 4:**
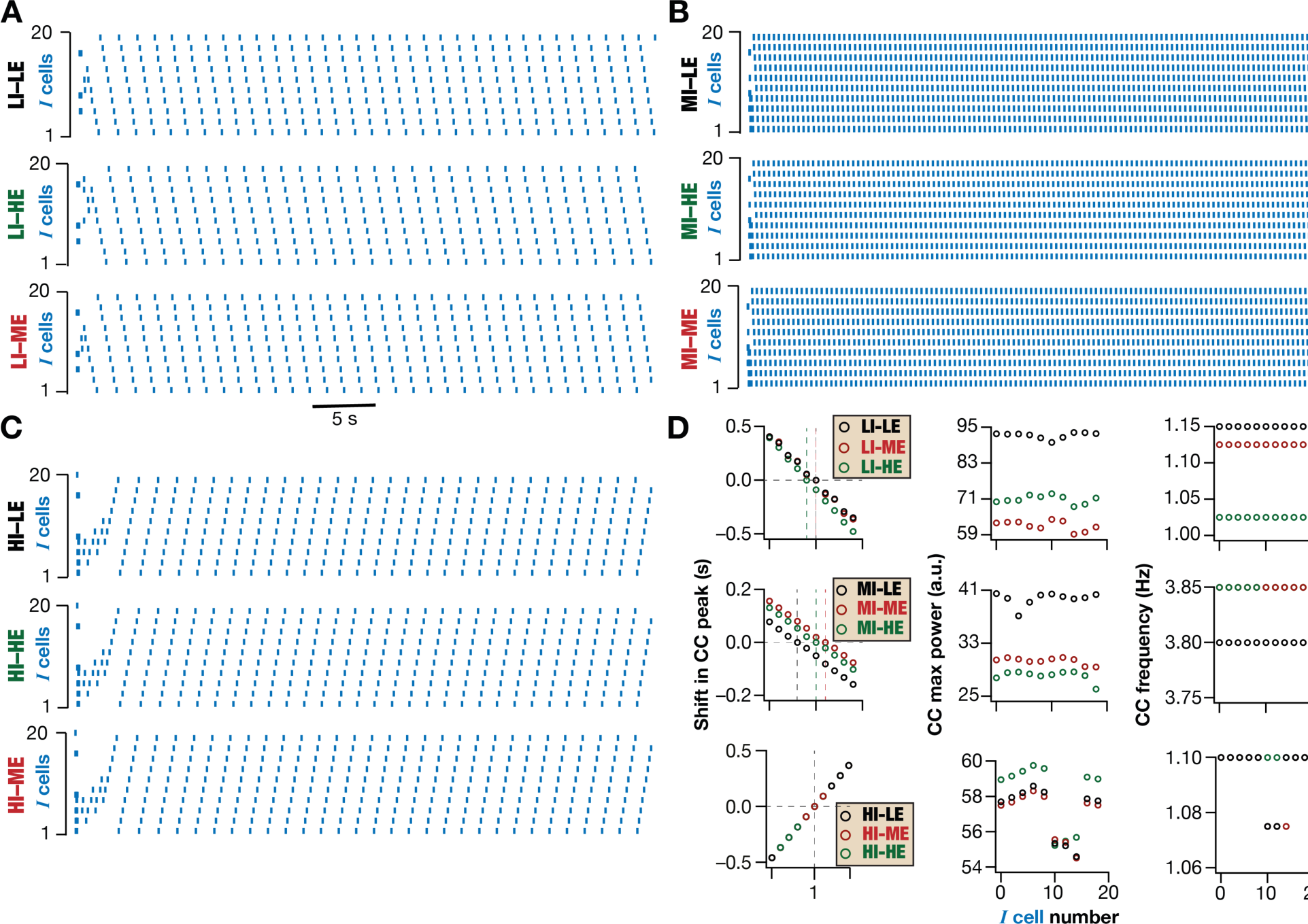
Robustness of patterned activity propagation in homogeneous ring networks constructed with E–I neurons with different excitability properties. Multiple conductance-based homogeneous E– I ring networks were built using distinct combinations of inhibitory and excitatory neurons. Neurons with low (L), medium (M), or high (H) excitability were chosen from the population of excitatory and inhibitory neurons. Nine homogeneous networks (80 *E* neurons, 20 *I* neurons) were constructed with 3 distinct *E* neurons and 3 distinct *I* neurons, with each subtype showing L, M, or H excitability. (A) Raster plots for the homogeneous ring network built with low excitability *I* neurons and low (LI–LE), medium (LI–ME), and high (LI–HE) excitability *E* neurons. (B) Raster plots for the homogeneous ring network built with medium excitability *I* neurons and low (MI–LE), medium (MI–ME), and high (MI–HE) excitability *E* neurons. (C) Raster plots for the homogeneous ring network built with high excitability *I* neurons and low (HI–LE), medium (HI–ME), and high (HI–HE) excitability *E* neurons. (D) *Left,* Plot showing the time points at which the cross-correlograms (with reference to *I* Cell #1, marked with color-coded dotted lines) peaked are shown for each *I* cell. The cells are arranged to depict the patterned propagation of the activity bump across the ring network. *Center*, Maximum power from spectral analysis of cross-correlograms of *I* Cell #1 with all the other cells. *Right*, Frequency of patterned activity computed from spectral analysis of cross-correlograms of *I* Cell #1 with all the other cells.

All 9 networks were tunable to exhibit stable propagation of patterned activity over the rings (Fig. 4) suggesting that these homogeneous ring networks could be realized with different sets of neurons with varied excitability to sustain patterned activity propagation. In these cases, we had hand-tuned the constant bias current onto the *I* neurons (Eq. 3) as well as the constant current required to elicit action potentials in *E* neurons towards achieving patterned activity propagation. Thus, the intrinsic excitability (*IE*) of the neuron needed to be in balance with the overall synaptic drive that is defined by the *E–I* balance of the circuit for robust function, further emphasizing the need to assess *E–I–IE* balance in networks rather than focusing on *E–I* balance (Seenivasan and Narayanan, 2020, 2022). We tested the robustness of these 9 hand-tuned homogeneous ring networks to noise introduced by high background activity. We used balanced high conductance state using several excitatory and inhibitory synapses impinging on each neuron as a source of synaptic noise (Fig. 4–figure supplement 1*A*). Without any additional tuning to the original 9 networks shown in Fig. 4, we tested their ability to propagate patterned activity across the rings in the presence of synaptic noise. We found the three ring networks with medium-excitability *I* neurons (MI–LE, MI–ME, MI–HE) to be the most stable to synaptic perturbations (Fig. 4–figure supplement 1*B*).

Together, our conductance-based homogeneous *E–I* ring network was scalable and could be constructed with neurons showing different values of intrinsic excitability towards manifesting stable activity propagation across the rings.

### Network-scale synaptic degeneracy in the manifestation of patterned activity propagation in homogeneous ring networks

As mentioned above, we had hand-tuned synaptic values to obtain a network that robustly sustained patterned propagation (Fig. 3). How sensitive was network function to the hand-tuned synaptic weight values? Were these synaptic weight values unique for the defined network towards sustenance of patterned propagation? Were there cross dependencies across synaptic weight values in the network required for sustaining robust propagation of patterned activity across the rings? To address these questions, we picked the homogeneous network in Fig. 3 and fixed the specific *E* and *I* neuron that we chose to construct the two homogeneous rings. We performed multiparametric stochastic search on the parameters governing synaptic weights in this homogeneous ring network towards assessing the sensitivity of patterned activity propagation to this synaptic parametric space. There were four parameters in our network that defined synaptic connectivity: the two weighting factors associated (one each) with the *I*-to-*I* and *I*-to-*E* Mexican-hat connectivity, the synaptic weight value assigned to the all-to-all *E*-to-*I* connections, and the parameter *α* in Eq. 3 that defined the coupling of neurons with afferent activity. To assess global sensitivity spanning all these parameters defining synaptic connectivity, we picked each of these four parameters from independent uniform distributions that spanned 0.5× to 2× of their respective base values. We constructed a ring network from each such randomized unique combination of these 4 parameters and repeated the network simulation towards generating network activity. A total of 500 unique ring networks were constructed through this process and activity patterns across the rings were recorded for each network for 36200 ms.

We found that while certain networks showed robust propagation of patterned activity, others failed to sustain patterned activity propagation (Fig. 5*A*, Fig. 5–figure supplement 1*A*). We used the spike timings recorded for all neurons in each of these networks to quantitatively identify those that sustained patterned activity propagation. Specifically, for each of these 500 ring networks, we measured the 3 correlogram-based quantitative metrics (CC slope, CV of *P*^::^, and CV of *f*^::^) that we had defined earlier (Fig. 5*B–D* for *I* neurons; Fig. 5–figure supplement 1*B–D* for *E* neurons). We set threshold values on each of these measurements towards achieving high CC slope and low values of both CVs. We declared those network models that satisfied these threshold values for all three measurements to be valid networks that showed patterned activity propagation (Fig. 5*A–D*, Fig. 5–figure supplement 1*A –D*). This validation process yielded a total of 327 valid ring networks (∼65%) that sustained patterned activity propagation along the ring.

**Figure 5:**
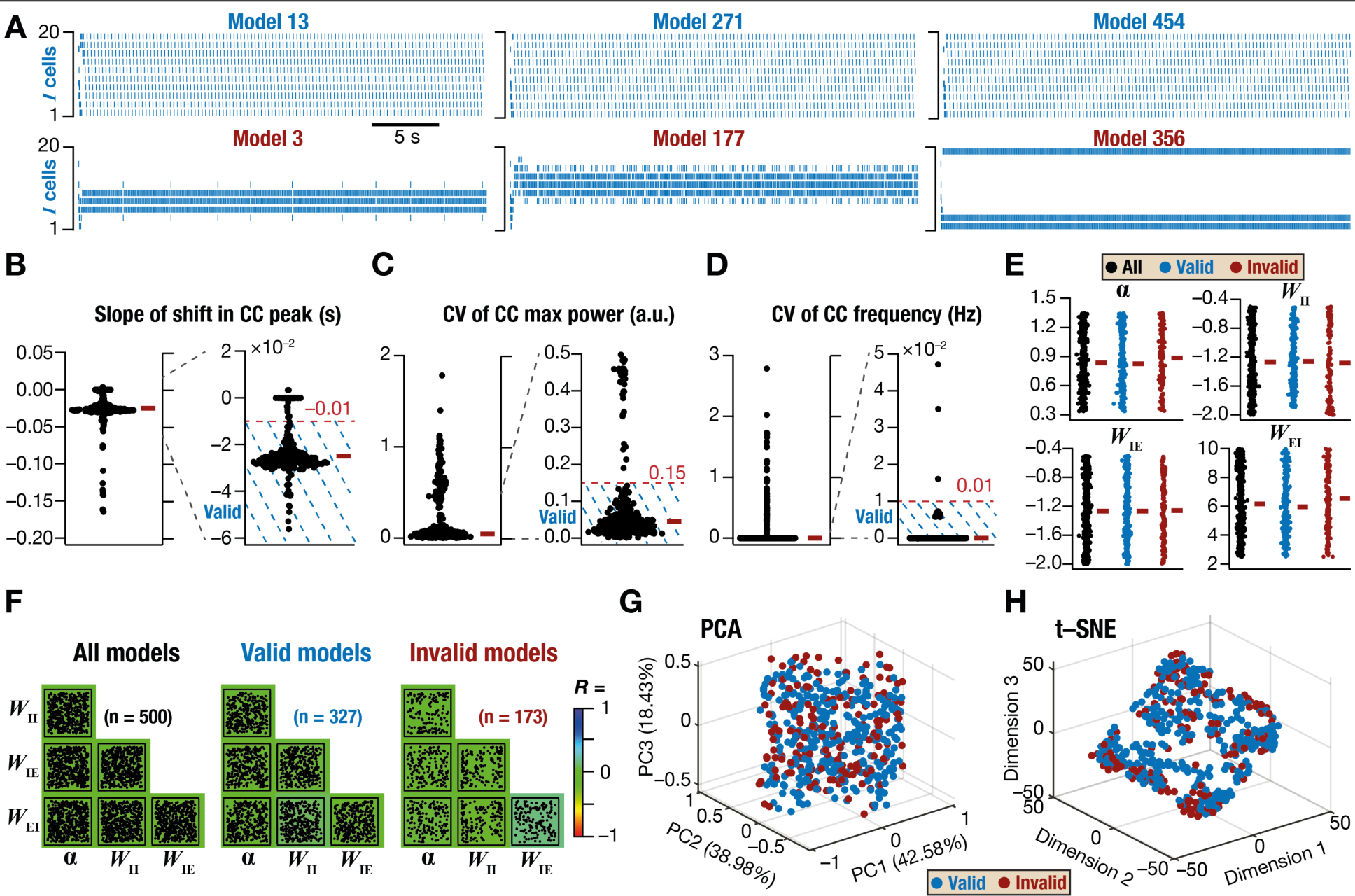
Sensitivity analysis of homogeneous ring network models revealed multiple synaptic routes to achieve robust propagation of patterned activity across the network. (A) A total of 500 homogeneous ring network models were generated by randomly altering the afferent (*α*) and local synaptic weight values (*W*IE, *W*EI, and *W*II) for the constructed network. Raster plots of example *I* neurons in network models exhibiting robust activity propagation (*Top*) and those that failed to exhibit patterned activity propagation (*Bottom*). The activity of example *E* neurons in these chosen models are shown in Fig. 5–figure supplement 1. (B–D) Validation of robust activity propagation was performed by employing thresholds on three metrics. Linear slope of the shift in peak location cross-correlogram (CC) between *I* Cell #1 and all other *I* cells in the network (*n* = 500 ring networks: 5 trials each on 100 networks). A threshold on the slope ensured that networks showing progressive shifts in the location of the activity bump were declared valid (B). A threshold requiring a low value for the coefficient of variance (CV) of the maximum power of the cross-correlogram (C), computed across all *I* cells, was used to declare models with high variability in maximal power across cells to be invalid. A threshold requiring a low value for the CV of the peak frequency of the cross-correlogram (D), computed across all *I* cells, was used to declare models with high variability in maximal frequency across cells to be invalid. While the non-zero slope of the shift in CC peak suggests attractor movement across the neural lattice, the lower CV for both maximum power and frequency implies similar firing properties across all cells in the network. Together, these three validation criteria were used to pick networks showing robust propagation of patterned activity. The red rectangle adjacent to each plot represents the respective median value. (E) Ranges of the 4 network parameters plotted for all (*n* = 500), valid (*n* = 327), and invalid (*n* = 173) models. The red rectangle adjacent to each plot represents the respective median value. It may be noted that all three distributions spanned a wide range of the sampled range in valid and invalid network populations. (F) Pair-wise correlations between 4 sampled network parameters for all, valid, and invalid models with the overlayed heat map of the correlation coefficient for each pair of parameters. It may be noted that the pair-wise correlation coefficients were weak for valid and invalid network populations. (G) Principal component analysis on the parameters associated with all valid (*n* = 327) and invalid (*n* = 173) networks. The variance explained by each dimension is shown within brackets. (H) *t*-SNE on the parameters associated with all valid (*n* = 327) and invalid (*n* = 173) networks.

How tightly clustered were the parameters that yielded the 327 valid ring networks? We plotted each of the four parameters that were used in the stochastic search process for valid and invalid networks and found them to span their respective ranges of permitted values (Fig. 5*E*). These results suggested that there was no clustering of synaptic parameters towards achieving ring networks that sustained patterned activity propagation. We then computed pairwise correlations on these parameters to ask if there were cross-dependencies across these parameters defining synaptic connectivity. We found pairwise Pearson’s correlation coefficients computed across all pairs of parameters, for valid and invalid models, were uniformly weak (Fig. 5*F*). These observations suggested a lack of pairwise interdependencies among these parameters in yielding networks that sustained patterned activity propagation. Finally, to determine the presence of lower-dimensional structures where valid model parameters could be clustered within, we performed linear (Fig. 5*G*) and nonlinear (Fig. 5*H*) dimensionality reduction techniques on the 4-dimensional parametric space. We did not observe any discernible lower-dimensional structure within which the valid network parameters clustered (Fig. 5*G–H*). Although there are strong global constraints on valid model parameters, discerned by the presence of invalid models whose parameters similarly spanned the same parametric space, the parameter values of valid ring networks were not sufficiently constrained to manifest as clusters or as strong pairwise correlations (Fig. 5*E–H*). These observations, showing that disparate combinations of synaptic weight parameters could yield valid ring network models, point to the expression of network-scale synaptic degeneracy in the emergence of patterned activity propagation in these homogeneous networks.

Together, our analyses showed that the weights associated with synaptic connectivity need not be fixed at precise values for the manifestation of robust propagation of patterned activity in ring networks. The manifestation of network-scale degeneracy with reference to synaptic weight parameters provides flexibility in the emergence of robust propagation of patterned activity, enhancing the degrees of freedom available to the network in sustaining such activity patterns.

### Increase in HCN-channel density enhanced the robustness of patterned activity propagation in homogeneous *E–I* ring networks

In a previous study, we had demonstrated that the suppression of low-frequency activity resulted in stabilization of grid-cell networks that manifested patterned propagation of neural activity along a 2D grid (Mittal and Narayanan, 2021). Our inspiration for the design of a biologically relevant mechanism to suppress low-frequency activation and introduce resonance into individual neurons was the HCN channel, a channel that imparts resonance onto the neural membrane they express by suppressing low-frequency activity (Hutcheon and Yarom, 2000; Narayanan and Johnston, 2008; Mishra and Narayanan, 2023). The ability to suppress activity arises from the gating properties of the HCN channel, which shut down an inward current upon depolarization and activate the same upon hyperpolarization of the neural membrane. This constitutes a negative feedback loop mediated by HCN channels that suppresses neural activity irrespective of the depolarizing or hyperpolarizing direction of input activity (Mishra and Narayanan, 2023). The additional requirement for the manifestation of resonance is the slow kinetics of HCN channels, which allows for the negative feedback loop to act predominantly for low-frequency inputs. As the channel’s slow kinetics do not allow them to sufficiently activate or deactivate during high-frequency inputs, HCN channels do not significantly alter high-frequency inputs. Thus, while a slow HCN channel would modulate excitability and introduce resonance, a faster HCN channel would still modulate excitability but would not sustain resonance (Narayanan and Johnston, 2008; Sinha and Narayanan, 2015) (Fig. 6*A*). Our earlier study had demonstrated that the slow kinetics of the negative feedback loop was an important requirement in its ability to stabilizing the CAN dynamics (Mittal and Narayanan, 2021).

**Figure 6:**
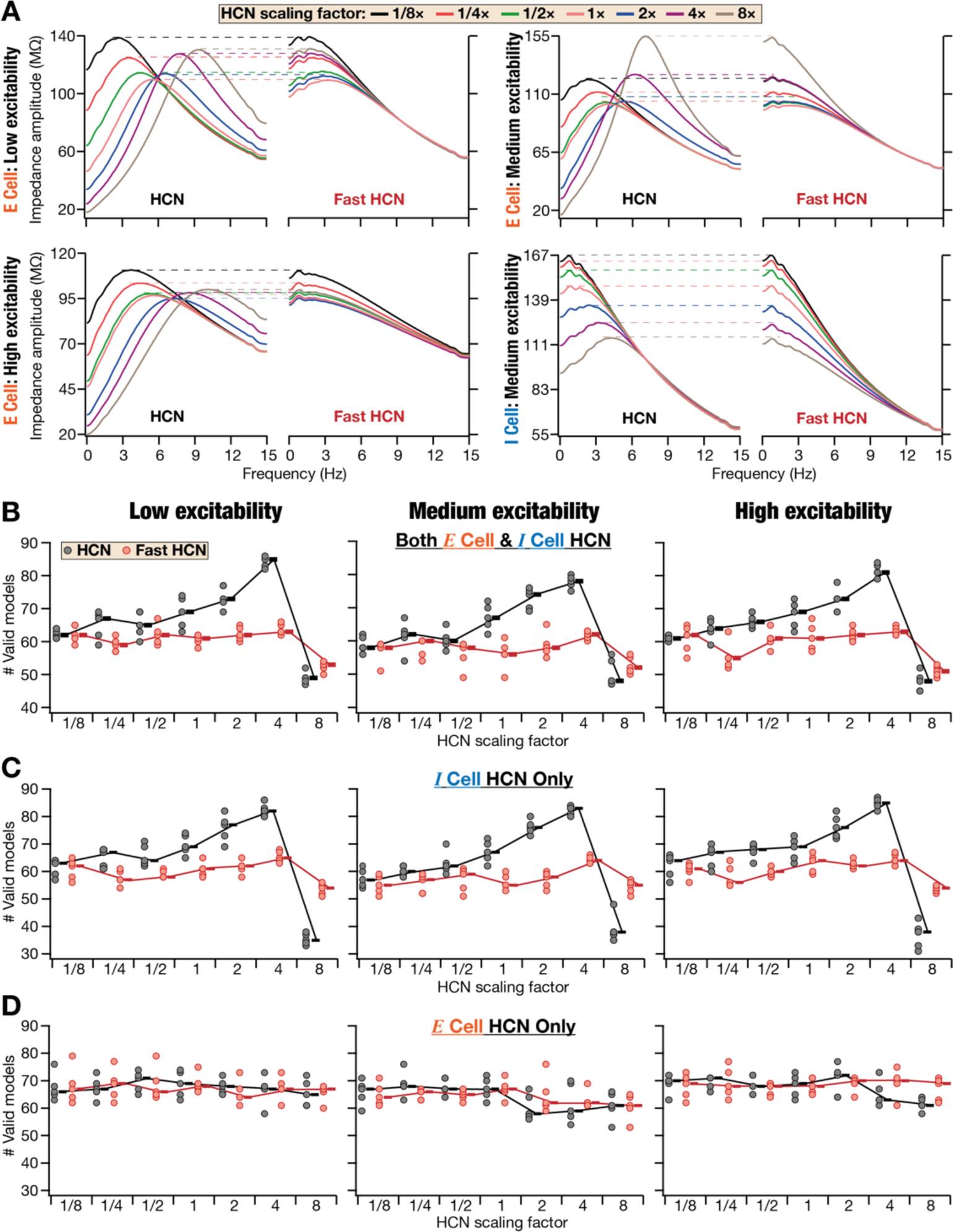
HCN channels stabilize patterned activity propagation in homogeneous ring networks. (A) Impedance profile and excitability of neurons with HCN and Fast HCN channels. The plots depict the procedure employed for matching the maximum impedance in the presence of HCN or Fast HCN channels. Examples are shown for how such matching is performed for *E* cells of different excitability and an *I* cell with medium excitability. Note that increasing resonance frequency increases with increasing HCN in all scenarios. (B) Impact of altering HCN conductance values in both excitatory and inhibitory neurons on the number of valid ring networks showing robust activity propagation. Intrinsically, all *E* cells are low (*Left*), medium (*Center*), or high excitability (*Right*) while the *I* cells were medium excitability for all three cases. Shown are the number of ring networks manifesting valid activity propagation when neurons were constructed with HCN or Fast HCN channels at different folds of change in HCN conductance values. (C–D) Same as (B) but with fold-change in HCN conductance limited to inhibitory (C) or excitatory (D) neurons. The *E* cells in (C) and *I* cells in (D) had respective baseline values of HCN conductance. For (B–D), for each HCN conductance fold-change level, 500 models (5 trials each with 100 networks) were obtained using a stochastic search on 4 network parameters mentioned in Figure 5. The rectangle adjacent to each plot represents the respective median value.

Motivated by these observations, our goal here was to assess if modulation of HCN channels in our conductance-based homogeneous ring network would alter the robustness of patterned activity propagation. In doing this, we first assessed the impact of altering HCN-channel density on the cellular physiology of *E* and *I* neurons. Expectedly, altering HCN-channel conductances in *E* or *I* neurons with slow HCN channels increased resonance frequency and modulated excitability (Fig. 6*A*). The specific quantum of changes in excitability depends on several factors, including the depolarizing shift to RMP introduced by HCN channels, the slow negative feedback loop triggered by time-varying inputs, and importantly on interactions with other channels that are expressed in the specific neuron under consideration (Mishra and Narayanan, 2023). In neurons with fast HCN channels, however, there was no resonance that was mediated by HCN channels (note that *E* neurons were endowed with other resonating conductances as well). To match subthreshold excitability (|*Z*|_.,5_) of the same neuron with fast or slow HCN channels, we adjusted the conductance of the fast HCN channels such that there was a match in |*Z*|_max_ with the same neuron with fast or slow HCN channels (Fig. 6*A*). Our experimental design was to construct ring networks with *E/I* neurons endowed with fast or slow HCN channels, expressed at different densities. Ring networks constructed with neurons endowed with different HCN channels were a means to delineate the impact of the expression profile, gating properties, and the kinetics of HCN channels on patterned activity propagation in the ring-network.

We employed the multiparametric stochastic search approach (on synaptic parameters, as in Fig. 5) to look for valid models, but with different types (fast *vs*. slow) and densities of HCN channels as well as using *E* neurons with different excitabilities (low, medium, and high). For each level of excitability, we performed 5 independent trials of network simulations (each spanning the 36200 ms period) on 100 different randomly generated network models (total 5×100 networks). We computed the cross-correlogram based metrics for each network and identified valid models that sustained robust propagation of patterned activity. We plotted the number of valid models for each trial made of 100 network models, together providing 5 numbers between 0–100 corresponding to the 5 trials. We then repeated these 500 trials (and all associated analyses) after increasing or decreasing HCN channels in *E* (5×100=500) or in *I* cells (5×100=500) or in *E* & *I* cells together (5×100=500), to 6 other HCN scaling factors (other than baseline). Our experimental design resulted in a total of 9500 (baseline 500 and 3×500×6 different HCN channel densities in different neurons) simulations with slow HCN channels for each level of *E*-neuron excitability. Finally, three distinct levels of *E*-cell excitability meant a total of 28500 (3×9500) simulations with slow HCN channels. A similar number of simulations were performed by replacing slow HCN channels by fast HCN channels, with everything else remaining the same. This process yielded a total of 57000 (2×28500) networks with fast/slow HCN channels with different densities in different neuronal subtypes with disparate levels of *E*-neuron excitability. The number of valid models (that showed patterned activity propagation) obtained for each trial was plotted against the density of HCN channels for each of the different configurations (Fig. 6*B–D*).

Under baseline conditions with medium excitability neurons (1× HCN scaling factor, slow HCN channels), we had earlier obtained ∼65% valid models (Fig. 5). Strikingly, with increase in HCN channel density (from 1/8× up to 4× of baseline levels) of both *E* and *I* neurons, there was a progressive increase in the number of valid ring networks that showed patterned activity propagation (Fig. 6*B*). These observations held irrespective of whether low, medium, or high excitability *E* neurons were used in building these networks (Fig. 6*B*). However, with very high HCN channel levels (8×), the excitability of neurons was reduced drastically, leading to fewer spikes in most networks. The reduction in spiking resulted in a sharp decrease in the number of stable networks when HCN-channel density was very high (Fig. 6*B*), thus providing an upper bound on the magnitude of increase in HCN channel densities towards stabilizing these networks.

The fast HCN channels (implemented by reducing their activation time constants, without altering voltage-dependence) are an effective way to delineate the impact of the contributions of excitability *vs*. dynamics of these channels. When we replaced all HCN channels (in *E* and *I* neurons) by their fast counterparts, we found that the beneficiary role of increasing HCN channel density, towards stabilizing patterned activity propagation, was lost (Fig. 6*B*). These observations emphasized the critical role of the slow kinetics of HCN channels in mediating the stabilizing effect observed in patterned activity propagation. Together, our analyses involving HCN channels in both *E* and *I* neurons showed that the possible number of synaptic parametric combinations that yielded valid ring networks could be increased with increase in HCN-channel density. The stabilizing role of HCN channels in yielding networks with robust propagation of patterned activity had an upper bound on the conductance and was dependent on the slow kinetics of the channel.

### Increase in HCN-channel density of inhibitory neurons enhanced the robustness of patterned activity propagation in homogeneous *E–I* ring networks

In our analyses involving HCN channels thus far, we had altered HCN channel kinetics and properties in *both* excitatory and inhibitory neurons (Fig. 6*B*). Were the stabilizing effects of increased HCN-channel density due to increased density in the *E* neurons or the *I* neurons? To address this, we assessed the outcomes of ring network simulations obtained with HCN-channel properties altered exclusively in all *I* neurons (Fig. 6*C*) *or* in all *E* neurons (Fig. 6*D*) without affecting the other subpopulation or the other properties in any way. We computed measurements based on the cross-correlograms across neurons with different HCN-channel densities and properties (Fig. 6–figure supplement 1) and found the number of valid models for each scenario (Fig. 6*C–D*). We found that increase in HCN-channel density exclusively in *I* neurons (Fig. 6*C*) mimicked the stabilizing effect observed with changes in both neural populations (Fig. 6*B*). Upon replacement of HCN-channels by their faster counterparts in *I* neurons, we found that the stabilizing role of HCN channels in sustaining patterned activity propagation was lost (Fig. 6*C*; Fig. 6–figure supplement 1). These observations held for networks constructed with *I* neurons with different levels of excitability (Fig. 6*C*). In contrast, limiting changes in HCN-channel density to the *E* neuron population did not provide any benefits in terms of increasing the number of valid networks that sustained patterned activity propagation (Fig. 6*D*). Consequently, there was no changes in the number of valid models obtained with either the slow or the fast HCN channels (Fig. 6*D*). These observations held for networks constructed with *E* neurons with different levels of excitability (Fig. 6*D*).

Together, these observations demonstrated increase in HCN-channel density in *I* neurons, but not in *E* neurons, enhanced the robustness of patterned activity propagation in the *E–I* ring network. The stabilization effect was dependent on the slow kinetics of HCN channels, as HCN channels endowed with fast kinetics were unable to enhance the number of valid models with increase in channel density. We also noted such specificity of the impact of HCN channels in the *I* neurons to be consistent with the design of the ring network where the propagation originated in the *I* ring owing to connectivity patterns spanning the *E–I* rings (Fig. 3*A*).

### HCN channels stabilize *E–I* ring networks with synaptic heterogeneities

Our analyses involving searches of synaptic parameters to identify valid networks (Fig. 5–6) did not account for heterogeneities in synaptic connectivity *within* the network. Although the synaptic weight parameters were altered, there were repeating in the same fashion for all neurons in the network. There was no difference in connectivity patterns between different *I* neurons or different *E* neurons. However, heterogeneities in parameters and measurements are ubiquitous across all scales of biological systems, and it is unrealistic to consider synaptic connectivity to be constructed with precise weights for all neurons in the network. How does the conductance-based *E–I* ring network behave with the introduction of heterogeneities in synaptic connectivity? Does the stabilizing role of HCN channels in homogeneous networks extend to networks with such synaptic heterogeneities?

We introduced synaptic heterogeneity into our conductance-based ring networks by introducing randomized additive jitter to the Mexican-hat connectivity defining the *I*-to-*I* connections (Fig. 7*A*). Except for this heterogeneity introduced in the connectivity pattern in the *I* ring, the network continued to be homogeneous with all *I* and *E* neurons being replicates of a single *I* and *E* neuron, respectively. Similar to our experimental design with homogeneous networks (Fig. 6), we analyzed such synaptically heterogeneous ring networks built with either low, medium, or high excitability *E*-neurons and medium excitability *I*-neurons. We systematically increased additive jitter to the synaptic weight matrix (Fig. 7*B*) and assessed jitter at three different levels to analyze changes in network physiology to progressive changes in synaptic heterogeneities. We generated 5 trials each for 50 network models (5×50=250) for the three degrees of synaptic heterogeneity (3×250=750) and three levels of *E*-neuron excitability (3×750=2250), yielding a total of 2250 network simulations (Fig. 7*C–E*). As before, these 2250 networks were analyzed with 6 other levels of slow HCN channel densities, providing a total of 15750 (2250×7=15750) simulations with different heterogeneities, different *E*-neuronal excitability values, and different HCN-channel densities. Finally, to assess the impact of slow *vs*. fast HCN channels, these 15750 simulates were repeated with fast HCN channels instead of the slow ones, yielding a total of 31500 networks over which the synaptic heterogeneity simulations were performed (Fig. 7). We computed the three correlogram-based measurements for each network and plotted the number of valid networks in each of the 5 trials as a function of HCN-channel density (Fig. 7*C–E*).

**Figure 7:**
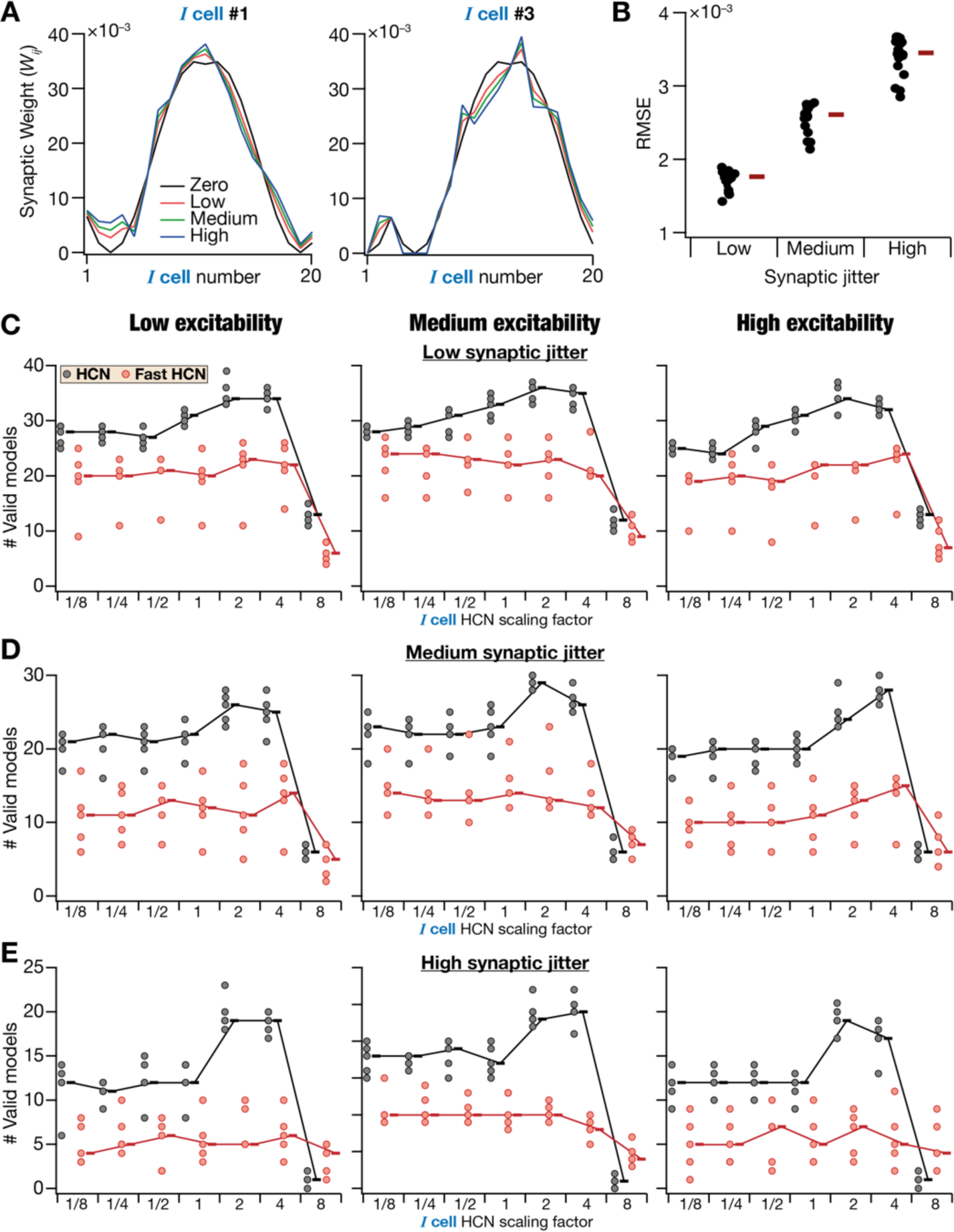
HCN channels stabilize patterned activity propagation in synaptically heterogeneous ring networks. (A) Synaptic weights depicting Mexican-hat connectivity of one of the *I* cells to all the other *I* cells in the network, in the absence or presence of increasing degrees of jitter. The Zero-jitter network will have identical synaptic weights for the connectivity of each cell to others (compare Zero jitter graphs for *I* Cell #1 (*left*) and *I* Cell #3 (*right*). On the other hand, when jitter is introduced, the connectivity patterns for each cell will vary from the other, thus introducing synaptic heterogeneities. Note that the weight values are distinct for *I* Cell #1 compared to *I* Cell #3. Also note that for a given instantiation, the low, medium, and high jitter conditions change in amplitude at each location, but their direction of change remains the same. Networks are run for different trials with different instantiations for each level of jitter (50 network models in each trial, with 5 independent trials). (B) Root mean square error (RMSE) was calculated between the synaptic weights under the “Zero jitter” case and synaptic weights for low, medium, and high levels of jitter, spanning all cells in the network. (C) Impact of altering HCN conductance values in inhibitory neurons on the number of valid ring networks showing robust activity propagation. Shown are the number of ring networks endowed with low synaptic jitter manifesting valid activity propagation when neurons were constructed with HCN or Fast HCN channels at different folds of change in HCN conductance values. (D–E) Same as (C) but for medium (D) and high (E) degrees of synaptic jitter. The rectangle adjacent to each plot represents the respective median value.

With increased degrees of synaptic heterogeneity, we observed a reduction in the number of ring networks that sustained patterned activity propagation (Fig. 7*C–E*). Specifically, with baseline level (1×) of slow HCN channels and low synaptic jitter, the percentage of ring networks that sustained patterned activity propagation was in the range of 60–65% (Fig. 7*C*). However, with increased synaptic jitter, this number reduced to ∼40–45% with medium synaptic jitter (Fig. 7*D*) and to 10–15% with high synaptic jitter (Fig. 7*E*). These results were consistent across networks with different *E*-neuron excitability (Fig. 7*C–E*). Thus, similar to their rate-based 2D counterparts (Mittal and Narayanan, 2021), synaptic heterogeneities destabilized patterned activity propagation in conductance-based 1D ring networks as well (Fig. 7*C–E*).

Strikingly, even in the presence of synaptic heterogeneities, increase in HCN-channel density in *I* neurons (upto 4×) resulted in an increase in the number of networks that sustained patterned activity propagation, irrespective of the level of excitability or the degree of jitter (Fig. 7*C–E*). Importantly, the beneficiary role of HCN channels in stabilizing patterned activity propagation was lost when HCN channels in *I* neurons were replaced by their fast counterparts. The numbers of valid models observed in networks with fast HCN channels were consistently lower compared to networks with slow HCN channels, across all excitability levels and all degrees of jitter (Fig. 7*C–E*). However, changes to HCN-channel density of *E* neurons did not yield a stabilizing influence on robust propagation of patterned activity (Fig. 7–figure supplement 1).

Together, these results demonstrated that the deleterious effects of synaptic heterogeneities on patterned activity propagation in the ring network could be suppressed by HCN channels in the *I* neurons. These results emphasized the critical role of the slow kinetics of the negative feedback loop (mediated by HCN channels) in conferring robustness of patterned activity propagation in synaptically heterogeneous ring networks.

### Intrinsic heterogeneities in inhibitory neurons destabilized patterned propagation of activity across ring networks

Neurons of any subtype manifest considerable heterogeneities in their intrinsic physiological properties, including excitability. Thus, the ring networks constructed thus far, involving repeating homogeneous *E* and *I* neurons within each ring, are biologically unrealistic. How does the manifestation of intrinsic heterogeneities in the neuronal population affect propagation of patterned activity in our ring network? To address this, we constructed ring networks with unique *E* and *I* neurons picked from their respective heterogeneous populations and assessed patterned activity propagation in these networks. Specifically, we generated 2000 ring networks where each inhibitory neuron (*n* = 20) and each excitatory neuron (*n* = 80) were unique, without altering synaptic connectivity. The *I* and *E* neurons were randomly picked from their respective heterogeneous populations of cortical interneurons (*n* = 930) and MEC LII stellate cells (*n* = 449). Surprisingly, none of these 2000 models exhibited stable propagation of patterned activity, demonstrating that intrinsic heterogeneities in the ring network have a strong deleterious impact on patterned activity propagation in the ring.

Was it heterogeneities in the *I* ring or the *E* ring that predominantly affected patterned activity propagation? To address this, we first constructed 100 ring networks where all *I* neurons were identical, but each *E* neuron was randomly picked from the heterogeneous stellate cell population. We observed that all 100 ring networks exhibited stable propagation of patterned activity (Fig. 8–figure supplement 1). Intrinsic heterogeneities in the *E* ring translated to heterogeneity in patterned activity (Fig. 8–figure supplement 1*A–D*) but did not disrupt patterned activity propagation across both rings (Fig. 8–figure supplement 1). We observed pronounced variability in the spiking activity of *E* neurons within and across different ring networks (Fig. 8– figure supplement 1*A*). However, such pronounced variability in spiking activity did not affect the validity of the network in terms of robustness of patterned activity propagation in *E* and *I* rings (Fig. 8–figure supplement 1*B–D*). We then limited intrinsic heterogeneities to the *I* ring of the network, letting all *E* neurons to be identical and by picking unique *I* neurons from the heterogeneous cortical interneuron population. Out of the 2000 randomized network models that we constructed by picking different sets of *I* neurons to construct these heterogeneous ring networks, none of them manifested robust propagation of patterned activity. Together, these observations demonstrated that intrinsic heterogeneities in the *I*-ring, but not the *E*-ring, had a strong deleterious impact on robustness of patterned activity propagation across the ring network.

**Figure 8:**
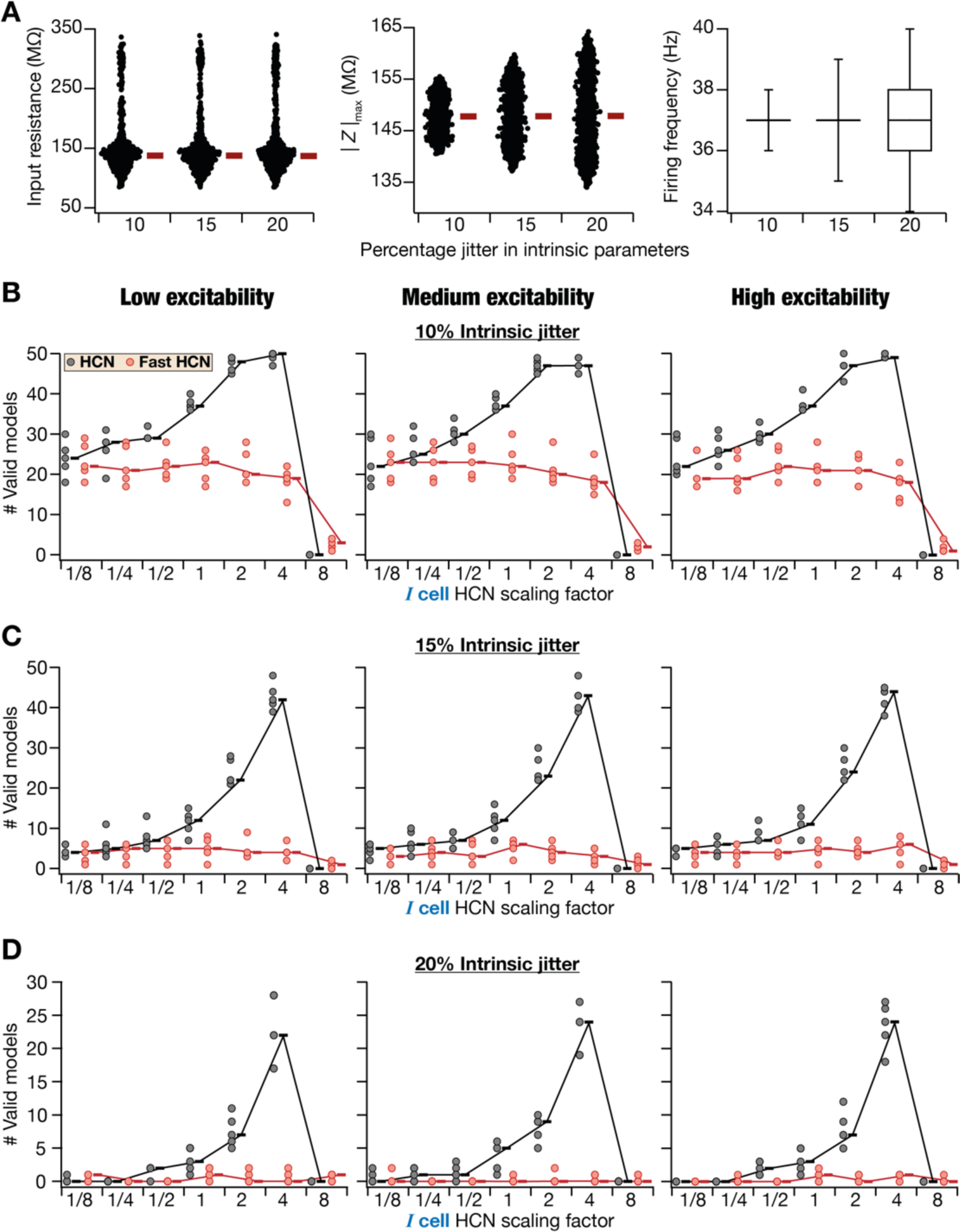
HCN channels stabilize patterned activity propagation in intrinsically heterogeneous ring networks. (A) Distribution of intrinsic measurements (input resistance (*R*in), impedance at resonance frequency (|*Z*|max), and firing frequency at 250 pA current injection (*f*250)) for different degrees of intrinsic heterogeneities introduced into inhibitory neurons. (B) Impact of altering HCN conductance values in inhibitory neurons on the number of valid ring networks showing robust activity propagation. Shown are the number of ring networks endowed with low intrinsic jitter manifesting valid activity propagation when neurons were constructed with HCN or Fast HCN channels at different folds of change in HCN conductance values. (C–D) Same as (B) but for medium (C) and high (D) degrees of intrinsic jitter. The rectangle adjacent to each plot represents the respective median value.

Quantitatively, how heterogeneous can the *I* layer neurons be for the ring network to be able to sustain patterned activity propagation? We picked *f*_250_, the firing rate of these neurons to a 250-pA current injection that varied between 30–47 Hz in the heterogeneous population of I neurons (Fig. 1*H*), as the representative measure of excitability. Instead of picking *I* neurons from this entire population of neurons, we picked different neurons that were endowed with the exact same *f*_250_ value of 37 Hz (the same *f*_250_ as the medium-excitability *I* neuron that we had used for the intrinsically homogeneous ring networks in Fig. 4–7). Although *f*_250_ was identical across these different *I* neurons, the density of the different ion channels and the other physiological measurements were heterogeneous (Fig. 8–figure supplement 2). Strikingly, ring networks built with *I*-neurons with the exact same value of *f*250, manifested robust propagation of activity across the rings, irrespective of the levels of excitability of neurons in the *E*-ring (Fig. 8–figure supplement 3). When the range of *f*_250_ was changed to 36–38 Hz (instead of a fixed 37 Hz) for picking randomized neurons, all networks failed to show patterned activity propagation without altering synaptic weights (specific hand-tuning was able to achieve activity propagation).

Together, these results demonstrated the deleterious impact of intrinsic heterogeneity, especially in the inhibitory layer of the network, on robust propagation of patterned activity across the ring network. Importantly, these results showed that equivalence of cellular-scale properties of individual neurons, despite heterogeneities in their underlying molecular mechanisms, sufficiently stabilized the network.

### HCN channels stabilized intrinsically heterogeneous CAN models

In introducing intrinsic heterogeneities, we had kept the synaptic connectivity constant and had incorporated intrinsic heterogeneities by different *I* neurons from the randomized population. Although these analyses showed that even small degrees of heterogeneities in suprathreshold excitability of *I* neurons can destabilize patterned activity propagation in the ring network, this form of introducing intrinsic heterogeneities was not conducive to assess the quantitative impact of different degrees of intrinsic heterogeneities on patterned activity propagation. Specifically, different neurons were built of different sets of ion channels and were endowed with variability in each of the different sub- and supra-threshold measurement. Thus, quantifying heterogeneities in different parameters and measurements and quantitatively linking them to loss of activity propagation becomes intractable given the number of parameters and measurements that governed the network. In addition, the analyses thus far did not explore potential routes to stabilize patterned activity propagation in a manner that was quantitatively demonstrable. To address these lacunae in our previous approach, we designed a second route to introduce intrinsic heterogeneities through perturbations to the base ring networks used earlier (Fig. 5–6). Specifically, we took the base network that showed patterned activity propagation and perturbed its intrinsic parameters by varying percentages and quantitatively assessed the network outcomes through correlogram-based measurements. The advantage with this approach was that the perturbations could be introduced at different degrees by adjusting the amount of jitter. This strategy provided a quantitative handle on the degree of intrinsic heterogeneities introduced and on whether there was a graded dependence of degradation (in patterned activity propagation) on intrinsic heterogeneities.

We added 3 distinct levels of jitter (10, 15, or 20 percentage jitter) to the baseline values of all channel parameters to yield heterogeneous *E*- and *I*-neurons, which were then used to construct heterogeneous ring networks (Fig. 8; Fig. 8–figure supplement 4; Fig. 8–figure supplement 5). Graded increase in intrinsic jitter of channel parameters expectedly yielded graded degrees of heterogeneities in intrinsic measurements of both *E* (Fig. 8–figure supplement 4) and *I* neurons (Fig. 8; Fig. 8–figure supplement 5*A*). We now employed the same strategy we had used for assessing the impact of synaptic jitter (Fig. 7) to quantitatively assess the impact of intrinsic jitter, HCN-channel density, and fast *vs*. slow HCN channels. A total of 31500 ring networks with 7 different densities of *I* cell HCN-channel, 3 different levels of E-cell excitability, fast or slow HCN channels, and 3 different levels of intrinsic jitter were analyzed. Each of the 5 trials for any given combination was made of 50 network simulations, whose outcomes were subjected to validation based on correlogram-based measurements. The number of valid models obtained in each trial was plotted against HCN-channel density for the different combinations (Fig. 8*B–D*).

With increase in intrinsic jitter, we observed a pronounced and progressive decrease in the number of valid ring networks that sustained patterned activity propagation (Fig. 8*B–D*). For instance, with medium-excitability neurons with baseline HCN conductance (1×) with slow kinetics, we obtained ∼70–80% valid ring networks with 10% intrinsic jitter (Fig. 8*B*), which dramatically reduced to ∼10–20% with 15% jitter (Fig. 8*C*), and further dropped to ∼5–15% with 20% jitter (Fig. 8*D*). These dramatic drops in networks that sustained patterned activity propagation were observed in networks constructed with any of the three levels of *E*-neuron excitability (Fig. 8*B–D*). We noted that the distributions of intrinsic properties of valid ring networks were comparable to their respective distributions across all models (both valid and invalid; Fig. 8–figure supplement 5*A*). Together, progressive increase in intrinsic heterogeneity of *I*-neuron excitability progressively and dramatically reduced the ability of the network to sustain patterned network activity.

Notably, the role of slow HCN-channels in stabilizing patterned activity propagation extended to networks with different levels of intrinsic heterogeneity as well (Fig. 8*B–D*). Specifically, increase in slow (but not fast) HCN-channel density in *I* neurons (upto 4×) resulted in an increase in the number of networks that sustained patterned activity propagation, irrespective of the level of *E*-neuron excitability or the degree of intrinsic heterogeneities (Fig. 8*B–D*). These results demonstrated the stabilizing role of *I*-cell slow HCN channels in intrinsically heterogeneous networks across a wide range of parametric combinations. However, there were limits to the ability of slow HCN channels in stabilizing intrinsically heterogeneous networks. First, as before, increasing HCN-channel density to 8× of baseline values led to significant drop in number of valid ring networks. Second, even at 4× increase in slow HCN-channel density, the number of valid ring networks progressively reduced with increase in intrinsic jitter. When we increased intrinsic heterogeneity to 30% intrinsic jitter, our simulations yielded no valid models under baseline values of slow HCN-channel density. However, there was a small increase in number of valid models with 4× increase in slow HCN-channel density (Fig. 8–figure supplement 5*B*).

Consistent with our previous analyses on E-ring intrinsic heterogeneities (Fig. 8–figure supplement 1), introducing heterogeneities through intrinsic jitter that was limited to *E*-neurons did not alter the number of valid ring networks (Fig. 8–figure supplement 5*C*). Furthermore, consistent with our results on synaptic heterogeneities (Fig. 7–figure supplement 1), changes to HCN-channel density of *E* neurons did not yield a stabilizing influence on robust propagation of patterned activity (Fig. 8–figure supplement 5*D*).

Together, these results demonstrated the critical role of the slow negative feedback loop mediated by *I*-ring HCN channels in stabilizing patterned activity propagation within intrinsically heterogeneous ring networks.

## DISCUSSION

The fundamental question that we addressed in this study was on how a phenomenon that emerged consequent to functional integration involving several functionally segregated subsystems was perturbed due to manifestation of heterogeneities. We considered heterogeneities either in the subsystems themselves or in the parameters that governed how the subsystems interacted with each other. The emergent phenomenon that we considered was patterned propagation of activity in a ring network that was constructed with several interacting neurons, each constituting an independent subsystem. While each subsystem (excitatory or inhibitory neuron) could be manifesting characteristic physiological properties, the underlying mechanisms (at the molecular scale: ion channels, pumps, buffers, etc.) that yielded these characteristic properties could be very different. Under such a scenario, does the emergent network-scale phenomenon depend on heterogeneities in cellular- or molecular-scale properties? Addressing this question demanded populations of neurons that showed similar cellular scale properties but were constructed with disparate molecular components (ion channels). Therefore, we constructed scalable conductance-based *E–I* ring networks that sustained patterned activity propagation across the rings with disparate sets of neurons and differential synaptic weight values. Importantly, these sets of neurons manifested characteristic physiological properties at the cellular scale, but were constructed with disparate sets of molecular components, thus allowing us to assess the impact of cellular-scale degeneracy on a network-scale phenomenon.

Our analyses demonstrated the deleterious impact of heterogeneities in neuronal or synaptic properties on patterned activity propagation in the ring network (Figs. 7–8). However, the impact of molecular-scale heterogeneities on patterned activity propagation was minimal when cellular-scale properties were matched. Importantly, HCN channels in the inhibitory neuronal population played a stabilizing role in sustaining patterned activity propagation in heterogeneous networks as well as in homogeneous networks endowed with synaptic heterogeneity. Using fast HCN channels, we demonstrated that this stabilizing impact of HCN channels on patterned activity propagation was critically reliant on their slow kinetics. The slow kinetics offer the means for targeted suppression of low-frequency signals and allow neurons to function as coincidence detectors for inputs arriving within a limited temporal window. Together, our results demonstrate that synaptic or intrinsic heterogeneities in ring networks were deleterious for patterned activity propagation but could be stabilized with the expression of HCN channels in the neurons that make the ring. In what follows, we explore the implications and limitations associated with our analyses and conclusions.

### Cascade of degeneracy spanning the molecular-cellular-network scales

A fundamental question in biological research is on how biological systems maintain functional robustness despite widespread heterogeneities in underlying components. Specifically, molecular components defining different neurons of the same subtype show widespread heterogeneities and neurons in a network manifest heterogeneities in their cellular properties. How does functional robustness emerge despite such widespread heterogeneities? A central answer that defines the ability of several biological systems to manifest robustness in the face of such heterogeneities comes in the form of degeneracy, the ability of disparate components to execute same or similar functions. When the same function can be executed using several combinations of different sets of underlying components, there is no necessity for maintaining homogeneous repeating subsystems. Different sets of functionally segregated subsystems can functionally integrate towards achieving the function, without compromising on the precision of the functional outcome. Degeneracy acts as a ubiquitous mechanism in complex biological systems to implement functional robustness by providing alternative routes in the face of biological variability, extrinsic perturbations, and loss of constitutive components (Foster et al., 1993; Edelman and Gally, 2001; Goldman et al., 2001; Prinz et al., 2004; Rathour and Narayanan, 2012, 2014; Mishra and Narayanan, 2019; Rathour and Narayanan, 2019; Goaillard and Marder, 2021).

Our study demonstrates a cascade of degeneracy (spanning the molecular, cellular, and network scales) in a ring attractor network whose function was assessed by its ability to patterned activity propagation, also assessing the impact of heterogeneities that manifest in systems that implement degeneracy. At the cellular scale, the different ion channels (molecules) constitute the functionally segregated subsystems with each ion channel endowed with distinct gating properties and kinetics. Our analyses demonstrated that disparate combinations of these distinct subsystems could functionally integrate to elicit characteristic cellular-scale physiological properties of neurons (Fig. 2). Despite dramatic differences in the set of ion channels expressed, their properties as well as the intrinsic neuronal properties of *E vs*. *I* neurons, this cellular-scale degeneracy was inherent to both the *E*- (Mittal and Narayanan, 2018) and *I*-cell (Fig. 2) populations. At the network scale, first, in homogeneous networks, our analyses showed that the weights associated with synaptic connectivity need not be fixed at precise values for the manifestation of robust propagation of patterned activity in ring networks. Several combinations of synaptic parameters were able to yield networks that sustained patterned activity (Fig. 5–6). In addition, when there was jitter induced in the synaptic connectivity patterns to make non-identical connectivity across different neurons, robustness of activity patterns was achieved by altering *I*-cell HCN channels (Fig. 7). The manifestation of network-scale degeneracy with reference to synaptic weight parameters provides flexibility in the emergence of robust propagation of patterned activity, enhancing the degrees of freedom available to the network in sustaining propagating activity patterns.

Importantly, network-scale degeneracy was not limited to synaptic parameters but extended to intrinsic properties of the neurons that were used to construct these networks. Specifically, activity propagation was robust when the *E–I* ring networks were constructed with heterogeneous *E* neurons and homogeneous *I* neurons (Fig. 8–figure supplement 1). More generally, activity propagation along the rings manifested lesser sensitivity to *E*-ring parameters compared to *I*-ring parameters (Fig. 6, Fig. 7–figure supplement 1, Fig. 8–figure supplements 1– 5). This is to be expected, given the architecture of the *E–I* rings whereby activity propagation was mediated by the *I* ring and was transmitted to the *E* ring through the Mexican-hat connectivity from the *I* to the *E* ring (Fig. 3*A*). Furthermore, activity propagation in the network was robust when heterogeneous models were chosen such that one excitability measure (firing rate for 250 pA current) was identical for all *I*-cells (Fig. 8–figure supplements 2–3). The underlying heterogeneities in other measurements and parameters of these models showed that robust activity propagation could be sustained without altering synaptic weights if the excitability of individual cells remained the same (Fig. 8–figure supplements 2–3). The heterogeneities in individual *I* cells parameters (Fig. 8–figure supplement 2) implied that disparate intrinsic components could yield similar activity propagation in the ring network (achieved with different *E*-cell excitability as well: Fig. 8–figure supplement 3). Finally, when all intrinsic measurements (including excitability) were forced to be heterogeneous through perturbations spanning all neurons, robust propagation was still achievable in several models by adjustments to HCN-channel densities in *I* cells (Fig. 8).

Together, our results demonstrate the manifestation of degeneracy in a network-scale function with respect to synaptic and intrinsic parameters that the network was built with (Fig. 5, Fig. 6, Fig. 7–figure supplement 1, Fig. 8, Fig. 8–figure supplements 1–5). Future analyses could focus on exploiting such degeneracy towards designing learning algorithms involving all network parameters (both synaptic and intrinsic) towards achieving robust activity propagation. Independent updates for all synaptic and intrinsic parameters towards achieving such goal will further enhance the possible combinations of parameters that could simultaneously yield robust activity propagation as well as characteristic physiological properties of the underlying cells.

### Physiological roles of HCN channels in the neuronal and network computations

In this study, we have shown that intrinsic neuronal resonance, which is mediated by HCN channels, imparts stability to the propagation of patterned activity across heterogeneous ring networks. Importantly, the slow kinetics of HCN channels played an important role in imparting this stabilizing role in the face of heterogeneities in and perturbations to synaptic and intrinsic properties of the neurons that constructed the ring network. The slow kinetics of HCN channels coupled to their voltage-dependent gating properties allow them to act as slow negative feedback loops that have been attributed to yield a stabilizing influence on the cells they express in (Hutcheon and Yarom, 2000; Biel et al., 2009; Zhao et al., 2009; Mittal and Narayanan, 2021; Mishra and Narayanan, 2023). This slow negative feedback loop allows for HCN-channels to act as biophysical mechanisms that suppress low frequency signals and 1/*f* noise (Hutcheon and Yarom, 2000; Narayanan and Johnston, 2007, 2008; Mittal and Narayanan, 2021). In addition, the slow negative feedback loop also equips neurons with the ability to switch across classes of excitability, reduce coincidence detection window (Das and Narayanan, 2014, 2015, 2017), and enhance spike phase coherence (Sinha and Narayanan, 2015, 2022).

Although the ability of HCN channels to regulate neuronal excitability and act as pacemaking channels have been studied extensively, this stabilizing role of HCN emanating from suppression of low-frequency signals and noise has not received commensurate attention. The relevance of somato-dendritic (Magee, 1998; Williams and Stuart, 2000; Lorincz et al., 2002) and dorso-ventral (Giocomo et al., 2007; Garden et al., 2008; Malik et al., 2016) gradients in HCN-channel expression as well as neuromodulation (Robinson and Siegelbaum, 2003; Biel et al., 2009) and activity-dependent plasticity (Fan et al., 2005; Brager and Johnston, 2007; Narayanan and Johnston, 2007, 2008; Mishra and Narayanan, 2022) in these channels to such stabilizing and low-frequency noise suppression should be systematically assessed (Mittal and Narayanan, 2021).

### Future directions

We built several heterogeneous conductance-based ring attractor networks that can sustain patterned activity propagation towards assessing the impact of heterogeneities in molecular and cellular components on continuous attractor networks. Our analyses provide fundamental clues about the manifestation of degenerate routes towards the emergence of continuous attractor networks and identify specific stabilization mechanisms that would improve robustness of continuous attractor dynamics in these networks. We had deliberately chosen a simple function (patterned activity propagation in 1D rings) to address these questions, given the complexities of the model and the parameters associated with each neuron in the *E–I* ring network. The fundamental insights gathered here about degeneracy (in terms of molecular and cellular components that build the CAN network) and stabilization mechanisms (slow feedback loop mediated by HCN channels) should be extended to complex CAN networks that execute other physiological roles (Khona and Fiete, 2022). In doing this, the specific architecture of the network, the intrinsic and synaptic properties associated with the different cells in the network, the type and pattern of external inputs they receive should all be accounted for, while explicitly factoring heterogeneities in each of these different components. Focus should also be on arriving at biologically plausible learning rules towards the emergence of such heterogeneous continuous attractor networks that manifest intrinsic and synaptic degeneracy, rather than merely identifying potential combinations of parameters that would yield such dynamics.

Future analyses could incorporate additional levels of biological details that would provide further insights about the multi-scale cascade of degeneracy and the stabilizing role of resonating conductances. To simplify computational complexity (which was already high owing to use of heterogeneous conductance-based neuronal models in networks demanding long network simulation time periods), we used single compartmental models for both excitatory and inhibitory neuronal subtypes. However, these neurons are endowed with elaborate dendritic arborizations and active-dendritic components, which play critical roles in single-neuron computation and how that translates to network outcomes (Johnston and Narayanan, 2008; Narayanan and Johnston, 2012; Major et al., 2013; Stuart and Spruston, 2015; Larkum, 2022). Therefore, there is a need to account for morphology and active dendrites into network models, but with detailed data on the ion-channel distribution and physiological properties along the somato-dendritic axis of these neurons. Such analyses should account for parametric variability of conductances, gating properties, and kinetics of all somato-dendritic ion channels rather than limiting the exploration to channel conductances (Rathour and Narayanan, 2014). In expanding analyses along these lines, it is essential to specifically identify the specific excitatory and inhibitory neuronal subtype involved in the function. Such delineation would enable explicit accounting of specific sets of constitutive components (*e.g.*, ion channels, receptors, buffers, and pumps), physiological measurements that define each neuronal subtype, and experimentally characterized heterogeneities in each component as well as in neuronal physiology.

Owing to computational limitations, our analyses was confined to one-dimensional (1D) continuous attractor model. Future studies could expand these analyses to two-dimensional (2D) conductance-based CAN models to assess degeneracy and stabilization in the emergence of for grid-patterned activity in these networks (Burak and Fiete, 2009; Mittal and Narayanan, 2021). It is essential to note that the computational complexity of such simulations is exorbitantly high with heterogeneous conductance-based models because the minimum network required for the emergence of grid-patterned activity is at least 40 × 40 = 1600 (compared to our 1D ring network with 20 *I* neurons; the *E*-ring merely follows the *I*-ring patterned activity). The orders of magnitude of increase in computational complexity involving different networks with varied heterogeneities and HCN-channel densities, each run for several trials, make 2D CAN networks difficult to be subjected to such elaborate conductance-based analyses. Conceptually, in performing such analyses, the specific subtype of excitatory (Domnisoru et al., 2013; Schmidt-Hieber and Häusser, 2013; Tang et al., 2014) and inhibitory (Rudy et al., 2011; Couey et al., 2013; Pastoll et al., 2013; Armstrong et al., 2016; Fuchs et al., 2016; Leitner et al., 2016; Tremblay et al., 2016) neurons, their connectivity patterns, intrinsic properties, and all associated heterogeneities need to be explicitly accounted for. Based on our analyses, we predict the manifestation of a cascade of degeneracy in the emergence of grid-patterned activity in 2D heterogeneous conductance-based CAN networks and a prominent stabilizing role for HCN channels in such emergence. These studies could also analyze the relative contributions of excitatory and inhibitory neurons to the emergence of attractor states (Burak and Fiete, 2009; Couey et al., 2013; Buetfering et al., 2014; Solanka et al., 2015; Miao et al., 2017; Winterer et al., 2017; Grosser et al., 2021; Tukker et al., 2022) in a conductance-based setting that explicitly accounts for all biological heterogeneities.

Future studies could focus on degenerate routes to achieving robustness of stable activity propagation. Our focus in this study was predominantly on the stabilizing roles of HCN channels. However, there could be several routes through which such stability could be achieved, given the different resonating conductances that express in several neuronal subtypes as well as the several routes to modulate resonance in these neurons (Hutcheon and Yarom, 2000; Hu et al., 2002; Narayanan and Johnston, 2007; Garden et al., 2008; Narayanan and Johnston, 2008; Hu et al., 2009; Zemankovics et al., 2010; Rathour and Narayanan, 2012, 2014; Rathour et al., 2016; Rathour and Narayanan, 2019). An extensive search process involving different ion channels and different synaptic connectivity could identify different mechanisms that could stabilize CAN networks. Importantly, all such future analyses mentioned in this section could have experimental counterparts, especially given developments in the large-scale recording techniques (Vyas et al., 2020; Steinmetz et al., 2021; Tukker et al., 2021; Gardner et al., 2022; Higley and Cardin, 2022; Urai et al., 2022; Zong et al., 2022) and neural-circuit manipulation techniques (Rajasethupathy et al., 2016; Kim et al., 2017a; Wolff and Olveczky, 2018; Nectow and Nestler, 2020; Shen et al., 2022) that target specific neuronal subsets.

Finally, conductance-based network models that explicitly account for molecular and cellular heterogeneities provide the ability to understand the implications for perturbing specific circuit components at the molecular or at the cellular scale. Therefore, they are amenable to addressing focused questions about how pathological conditions that alter specific molecules (ion channels, receptors) confined to certain neuronal subtypes affect continuous attractor networks and associated behavioral outcomes. As synaptopathies and channelopathies are strongly associated with several neurological disorders (Kullmann, 2002; Terzic and Perez-Terzic, 2010; Grant, 2012; Poolos and Johnston, 2012; Lerche et al., 2013; Brager and Johnston, 2014; Mandolesi et al., 2015; Rivolta et al., 2020; Deng and Klyachko, 2021; Franklin and Laubham, 2021; Kessi et al., 2021; Mantegazza et al., 2021), this ability to assess the *heterogeneous* impact of altering specific components is invaluable from a theoretical perspective as well as from the perspective of designing specific experiments or drug targets. The process outlined here to generate heterogeneous networks could also be employed to build heterogeneous neurons and networks that mimic the several differences that are observed under specific pathological conditions. Such analyses would enable deeper understanding of the impact of all the concomitant changes (Mishra and Narayanan, 2021) occurring with that pathological condition, rather than assessing impact using piece-meal analyses involving each component individually. Together, the procedures and the *heterogeneous* networks described in our study provide insights about the combinations of mechanisms that would sustain and stabilize continuous attractor dynamics in networks that drive a variety of behavioral goals (Khona and Fiete, 2022) under physiological and pathological conditions.

## METHODS

We constructed conductance-based ring network models with both excitatory (*E* cells) and inhibitory (*I* cells) neurons towards testing the robustness of patterned activity propagation across the ring in the presence of neural-circuit heterogeneities and stochasticity. To this end, the first task was to build heterogeneous populations of biophysical models for both inhibitory and excitatory cells. We employed stellate cells from the entorhinal cortex as representative excitatory neurons and used a conductance-based heterogeneous population of stellate cells from a previous study (Mittal and Narayanan, 2018). We built a conductance-based heterogeneous model population of cortical interneurons to act as representative inhibitory neurons. Homogeneous or heterogeneous ring networks were then constructed using these conductance-based excitatory and inhibitory neurons, with characteristic connectivity patterns associated with typical ring attractor networks. We defined specific metrics to assess patterned activity propagation in ring networks to assess robustness in the face of different forms of perturbations, heterogeneities, and noise. Finally, in scenarios where loss of patterned activity propagation in the ring network was observed, we assessed potential remedial mechanisms that could stabilize patterned activity propagation across the ring network. In what follows, we describe the methodological details associated with each of these steps.

### Heterogeneous populations of excitatory and inhibitory neurons

#### Heterogeneous population of excitatory neurons

We used single compartmental conductance-based models of stellate cells (SCs) from layer II (LII) of medial entorhinal cortex (MEC) with 9 active conductances and passive leak channels as excitatory neuronal models. The rationale behind the choice of LII SCs as excitatory neurons was that a proportion of these neurons act as grid cells, whose patterned activity has been effectively explained using the continuous attractor network (CAN) framework. As our goal was to construct a ring attractor network made of conductance-based models, we employed these neurons as representative excitatory neurons. We used a physiologically-validated (validated against 10 electrophysiological measurements from SCs) heterogeneous population of SCs from a previous study (Mittal and Narayanan, 2018). We used a subset of the total 449 valid SCs that were available (Mittal and Narayanan, 2018).

#### Heterogeneous population of inhibitory neurons

For inhibitory neurons, we built a population of single-compartmental conductance-based models of fast-spiking cortical basket cells. The single compartment model was in shape of a cylinder with diameter and length of 75 *μ*m. The choice of using a single compartment model by the need for a computationally simple model that can be part of a large network. The rationale behind the choice of cortical interneurons as inhibitory neurons was that the excitatory neurons were from the cortex and the availability of electrophysiological data for cortical interneurons.

#### Passive and active properties of cortical interneuron models

The passive properties of the interneuron models were specified by specific membrane resistance, *R*. (specified in kΩ.cm^2^) representing leak channels (*g*_;8,9_= 1/*R*.) and specific membrane capacitance, *C*. (fixed at 1 µF/cm^2^). We introduced 5 different voltage-dependent ion-channels into this model: fast sodium (NaF), delayed rectifier potassium (KDR), hyperpolarization-activated cyclic nucleotide-gated (HCN), *D*-type potassium (KD), and *A*-type potassium (KA) channels (Fig. 1*A*). All channels were modeled using the Hodgkin-Huxley formulation (Hodgkin and Huxley, 1952), with gating and kinetics properties for all the channels derived from a previous study (Tzilivaki et al., 2019). All channels followed the Ohmic formulation for their current-voltage relationships, with the reversal potentials of Na^+^, K^+^, and HCN set at +50, –90, and –10 mV, respectively. The reversal potential for leak channels was set at –70 mV. The resultant 6 active and passive parameters that were varied across models are listed in Table 1.

#### Intrinsic physiological measurements of cortical interneurons

A set of intrinsic physiological measurements were computed from interneuron models, towards characterizing and validating them. Resting membrane potential (*V*RMP) was measured after allowing the cell to settle for 5 s when no current was injected and was computed as the mean membrane potential for a 1-s period between 5–6 s (Fig. 1*B*). This was important as there were conductances in our cortical interneuron model that were active at rest and therefore contributed to resting membrane potential. The passive membrane potential, with just leak channels, was set at –70 mV and then the interactions among the resting conductances were allowed to guide the membrane potential towards the steady-state *V*RMP. To ensure that *V*RMP was stable and devoid of any spontaneous oscillations or spiking activity, we computed the standard deviation of the membrane potential between 5–6 s duration when no current was injected (Fig. 1*B*). We set a bound on the maximum possible standard deviation of membrane potential under resting conditions towards ensuring the absence of voltage fluctuations in valid models (Table 2). All the other intrinsic measurements detailed below were computed after the voltage settled at this *V*RMP, that is, at the 6-s time point after simulations started.

Sag ratio (*Sag*) was measured as the ratio of the steady-state membrane potential deflection (*V*SS) to peak membrane potential deflection (*V*peak) in the voltage response of the model to a hyperpolarizing step current of 200 pA for a duration of 1000 ms (Fig. 1*C*). We recorded the number of action potentials elicited by model neurons for depolarizing current pulses of 50 pA (*f*50; Fig. 1*D*) or 250 pA (*f*250; Fig. 1*E*) for a duration of 1s. We computed the input resistance (*R*in) as a measure of subthreshold excitability of model neurons by injecting 1-s long step currents ranging from –100 pA to 100 pA in steps of 20 pA and recorded the steady-state voltage responses (Fig. 1*F*). The steady-state voltage response was plotted against the corresponding amplitude of current injection and *R*in was measured as the slope to this *V*–*I* plot (Fig. 1*F*). Spike amplitude (*V*AP) was computed as the difference between the peak voltage of the action potential and *V*RMP (Fig. 1*G*). Threshold of action potential (*V*th) was defined by the membrane potential at which the first derivative of action potential crosses 20 V/s (Fig. 1*G*). Action potential half-width (*T*APHW) was measured as the temporal width of action potential at point where voltage was halfway between maximum amplitude achieved by action potential and the action potential threshold (*V*th) (Fig. 1*G*). Afterhyperpolarization (*AH*P) was measured as the difference in membrane potential between action potential threshold (*V*th) and the lowest value of membrane potential achieved after the action potential (Fig. 1*G*).

#### Multi-parametric multi-objective stochastic search (MPMOSS) algorithm

To generate a heterogeneous population of physiologically-validated cortical interneurons, we used the MPMOSS algorithm (Foster et al., 1993; Goldman et al., 2001; Prinz et al., 2003; Marder and Taylor, 2011; Rathour and Narayanan, 2012, 2014; Anirudhan and Narayanan, 2015; Srikanth and Narayanan, 2015; Mukunda and Narayanan, 2017) similar to the one that was used in arriving at the excitatory neuronal population (Mittal and Narayanan, 2018). First, a specific neuronal model was generated by independently picking a unique random value for each model parameter from their respective uniform distributions (Table 1). Several such randomized models were generated by repeating this process multiple times (10,000). Characteristic properties of each randomly generated model (Fig. 1) were computed and compared with their electrophysiological counterparts (Table 2). This stochastic search process was performed over 6 model parameters (Table 1) and the randomized neurons were validated against electrophysiological bounds of 10 sub- and supra-threshold intrinsic measurements (Table 2). The models that satisfied the bounds associated with all 10 measurements were declared as valid models. We obtained 930 valid models (out of the 10,000 generated; 9.3% valid models) of cortical interneurons at the end of this search process.

In assessing this model population, pairwise correlations between the 9 different intrinsic measurements (Table 2; *f*50 was not included in this analysis because *f*50 was required to be zero for all valid models) were computed to assess pairwise relationships between different intrinsic properties across the valid model population. Assessing these correlations provided us insights about whether the different measurements obtained from the model populations were indeed assessing different aspects of model behavior, or if they were just cross-dependent measurements constituting minimal cross-measurement constraints on the parametric space. To assess parametric interdependencies, we computed pairwise Pearson’s correlation coefficient between the 6 parameter values (Table 1) obtained from the different valid models (Fig. 2*H–I*).

### Structure and dynamics of the E–I ring network

Our *E–I* ring network model comprised of an inhibitory (*I*) and an excitatory (*E*) layer, with neurons arranged in a ring-like architecture within each layer (Fig. 3*A*). The *I:E* neurons ratio was maintained at 1:4, with the default size of the *I* and *E* layers set at 20 and 80, respectively. For homogeneous ring networks, all *I* neurons in the model were identical in terms of their intrinsic parameters and properties, picked from the valid model population of cortical interneurons. To explore the impact of excitability of *I* neurons, different sets of homogeneous networks were constructed such that the picked *I* neuron had low, medium, or high excitability. Similarly, all excitatory neurons were identical in terms of their intrinsic parameters and properties, picked from the valid model population of stellate cells. Three distinct *E* neurons with low, medium, and high excitability were picked to construct different sets of networks with different *E*-neuron excitability.

#### Inputs to neurons of inhibitory layer of the network

The inhibitory ring network was constructed as a one-dimensional equivalent of the 2-D continuous attractor network that generated grid-patterned network activity (Burak and Fiete, 2009). Similar to the 2D network (Burak and Fiete, 2009), each neuron in the *I*-ring was considered to receive a head-direction and a velocity signal, albeit along a virtual one-dimensional arena as opposed to a 2D arena in the grid-cell network. Specifically, each neuron in the *I*-ring had a direction preference *μ*_<_ (with an assumption that each neuron in the network was receiving specific head-direction inputs), which can be either π/2 or 3π/2 depicting left and right movement, respectively. Within the *I*-ring, left and right neurons were arranged in an alternate manner. We connected these *I*-neurons through a center-shifted Mexican-hat connectivity, towards implement biologically relevant connectivity that results in the patterned activity propagation (Burak and Fiete, 2009):

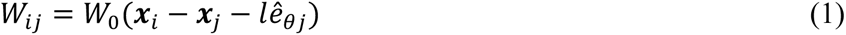

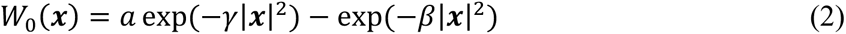

where, *w*_<=_ represented the synaptic weight from neuron *j* (located at *x*_=_) to neuron *i* (located at *x*_<_) within the *I*-ring, *e*^_>=_defined the unit vector pointing along the *θj* direction. For coupling afferent inputs with the propagation of activity within the *I*-ring, a center-shifted asymmetry was introduced in the Mexican-hat connectivity between *I* cells (*w*_II_) based on direction preference *μ*_=_of the neuron. The parameter *l* with a default value of 2 neurons defined the amount of shift along *e*^_>=_. This resulted in self-inhibition as the value of *a* in the difference of Gaussian formulation was set at 1, together defining all-inhibitory connectivity among *I* cells. All the inhibitory synaptic connections were made using a conductance-based GABAA receptor model. The other parameters were *γ* = 1.1×*β* with *β* = 3/λ^2^, and λ (default value: 13) defining the periodicity of the *I*-layer neural lattice (Burak and Fiete, 2009; Mittal and Narayanan, 2021).

The other set of inputs to the *I*-cells were the feed-forward inputs, which have information coming from outside of the network. If we consider our network to be a one-dimensional version of the grid-cell activity network, this feed-forward input would represent the velocity (*v*) of the traversing animals with modulation by the preferred direction of the neuron (*e*^_>=_). If we consider our network to be the angular path integration network for head-direction cells in mammals or ellipsoid body in drosophila, then the afferent input would represent the angular velocity with modulation by direction preference of the neuron. As our analyses was agnostic to the specific function of the network, we called these inputs as afferent inputs, and the current associated with this input was computed as:

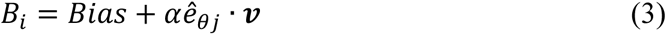

where *α* denotes a gain parameter for the afferent inputs (afferent inputs scaling factor), *v* represents the velocity or angular velocity vector derived from the 1D trajectory of the virtual animal. For each time step, the value of *v* was picked from a uniform distribution (0–4 m/s or s^−1^). In most simulations, the direction of velocity remained fixed along one direction. Bias was a constant depolarizing current adjusted according to intrinsic excitability of the cortical interneurons. The constant *Bias* current could be considered as a simplified abstraction of the overall excitatory activity impinging on these neurons, similar to the 2D version of the model (Burak and Fiete, 2009) that is adapted here to a 1D version.

The last set of inputs to neurons to the *I*-ring was driven by synaptic inputs from the excitatory layer. These synapses were driven by all-to-all connectivity from excitatory neurons to inhibitory neurons (*w*_@?_). All excitatory connections were modeled as conductance-based AMPA receptors. This architecture effectively contributes to the design of a conductance-based 1D continuous attractor network implementing the velocity shift mechanism (Burak and Fiete, 2009; Khona and Fiete, 2022).

#### Inputs to neurons of excitatory layer of the network

The *E* cells in the network were also arranged in a ring-like manner but have no direction preference and lack connectivity among themselves. The *E* ring was representative of how the *I*-ring cells connected to the *E* cells. Specifically, *E* cells received inhibitory inputs following the repetitive Mexican-hat connectivity (*w*_?@_) from the *I*-ring. They also receive constant depolarizing input that depended on the intrinsic excitability of the specific stellate cell model chosen to construct the network.

#### Excitatory and inhibitory synapses

The *E*-to-*I* connections were modelled using synapses containing AMPA receptors while *I-*to-*I* and *I*-to-*E* connections were modelled as synapses containing GABAA receptors. Both receptors followed a 2-state gating model that switched from closed to open state upon transmitter binding, and their currents were modeled using the Ohmic framework. The reversal for AMPA receptor was 0 mV and for GABAA receptor was –80 mV. The rise time and decay time for both receptors were 0.1 ms and 10 ms, respectively. Action potentials (APs) in the identified presynaptic neuron were detected as positive-going zero-crossings of the presynaptic membrane voltage. A synaptic transmission delay of 4 ms was introduced between the detection and introduction of a postsynaptic current in the identified postsynaptic neuron.

#### Balanced high conductance state (HCS)

The robustness of our network was tested in the presence of *in vivo*-like conditions where background synaptic activity was prevalent. To implement this, we introduced designated excitatory and inhibitory synapses (different from the ring-network synapses described above) into all neurons. Balanced high conductance state (HCS) was achieved by tuning the number (excitatory synapses: 100 and inhibitory synapses: 20), the conductance and average firing frequency (excitatory synapses: 3 Hz and inhibitory synapses: 10 Hz) of these synapses. The overall impact on the network was the manifestation of noisy fluctuations around resting membrane potential on the model neuron as a consequence of open synaptic receptors that yield no average net current. Both excitatory and inhibitory synapses were modeled using the formulation involving the sum of two exponentials, one for the rise (*τ*_ABC2_ = 2 ms) and the other for the decay (*τ*_D2E/F_= 10 ms) of the associated synaptic current. The reversal potential for excitatory synapses was 0 mV and for inhibitory synapses was –80 mV.

#### Network simulation details

For the stabilization of the resting membrane potentials of neurons in the network, no inputs were given for the initial 6 s period after simulations were started (*RMP stabilization period*). After this stabilization period of 6 s, each *I* cell was injected with a depolarizing step current whose amplitude was randomly picked from a uniform distribution spanning 0–200 pA and was injected for 100 ms to initialize the network dynamics (*network initialization*). For different trials of the simulation, the network initialization was randomized while keeping all the other network and neuronal parameters the same. As a final step in the initialization period, the velocity input was zero for the additional 100-ms period (*afferent input stabilization*) to allow an initial constant feed-forward drive during this period (*v* in Eq. 3 was set to zero during this 100 ms period). Beyond this initialization period, there was no additional current injection (the random initialization current was limited to a 100 ms period) and velocity inputs were allowed to continue as in Eq. 3. Together, the entire duration for the simulation was 36.2 s, with 6 s of *V*RMP stabilization, 100 ms of network initialization, 100 ms of afferent input stabilization, and 30 s of actual simulation to evaluate patterned activity propagation. All raster plots depict firing during these 30 s of actual simulation period. Of these 30 s, the initial 5 s was considered the transient period where patterned activity emerged and stabilized in propagation across the rings. Therefore, all analyses associated with patterned activity propagation were performed on the last 25 s of data (11.2–36.2 s). Patterned activity propagation was found to be dependent on network configurations as well as the specific randomization associated with the network initialization process.

In one set of simulations (Fig. 3–figure supplement 3), the direction of velocity inputs was abruptly flipped by 180° at the 10 s time-point after the end of the stabilization period to test if patterned activity propagation in the ring changed directions with change in sign of the velocity inputs.

### Sensitivity analysis on network parameters

We tested the robustness of our conductance-based ring network model for different combinations of four network parameters: afferent input scaling factor (*α* in Eq. 3) as well as scaling factors for the three different synaptic weight matrices associated with *I–I* (*w*_II_), *I–E* (*w*_?@_), and *E–I* (*w*_@?_) connectivity. Sensitivity analysis involving perturbation to all four network parameters was simultaneously assessed using an algorithm similar to the MPMOSS procedure employed in the generation of individual neuronal populations. However, here, the parameters over which analysis was performed defined deviations of these four network parameters from their default values. Specifically, sensitivity analysis was performed by randomized picking of each of these four network parameters from independent uniform distributions that spanned 0.5×–2× of their respective default values (perturbations from default values). A population of ring networks was construction by repeatedly sampling the independent distributions associated with the four network parameters. Simulations to assess patterned activity propagation across the network were performed on each such randomized network. The manifestation of robust patterned activity propagation was assessed using quantitative measurements derived from the activity of neurons in the network. Thus, the multiple parameters associated with this MPMOSS algorithm were the deviations from default values of the four network parameters, and the objectives together assessed whether the network manifested patterned activity propagation after introduction of these perturbations.

### Network heterogeneities

We introduced two forms of heterogeneities into our conductance-based ring network model, one by introducing heterogeneities to the synaptic connectivity matrix and another by altering the intrinsic properties of the different neurons in the network.

#### Synaptic heterogeneities

The neurons in the *I*-layer of the network were connected to each other following precise Mexican-hat connectivity (Eq. 1). To account for biological heterogeneities, including presynaptic properties, postsynaptic receptor densities, bouton and spine geometry, dendritic processing, and local plasticity mechanisms, that may affect such precision connectivity, we introduced additive random jitter to the Mexican-hat connectivity that defined the *I*-to-*I* synaptic weights *w*_II_. We achieved three degrees of synaptic heterogeneities (low, medium, and high) by adjusting the extent of jitter that was introduced.

#### Intrinsic heterogeneities

An intrinsically heterogeneous network was defined as a network where each neuron exhibited distinct values for their intrinsic parameters (passive and active components) as well as their characteristic electrophysiological measurements (sub- and supra-threshold properties). We used two distinct strategies for the incorporation of intrinsic heterogeneities into the network. In the first strategy, we used the heterogeneous population of valid *E* and *I* cells generated using the respective MPMOSS algorithms to pick unique neurons representing each *E* and *I* cell in the network. Specifically, for the *I*-layer, we randomly picked 20 distinct model neurons from the 930 valid cortical interneurons. For the *E*-layer, 80 intrinsically heterogeneous model neurons were randomly picked from the 449 valid MEC LII stellate cells. Thus, the resultant network model exhibited intrinsic heterogeneity in both *E*- and *I*-layers of the network. We repeated this process 2000 times (different sets of 20 *I* cells and 80 *E* cells were randomly picked each time) to construct 2000 distinct intrinsically heterogeneous networks. When heterogeneity was to be limited to one of the two layers (*E*- or *I*- layer), the process was similar for the heterogenous layer with the neurons in the other layer retained to be homogeneous.

The second strategy we used to incorporate intrinsic heterogeneity was by introducing random additive jitter to the channel parameters (all channel conductances and specific membrane resistance; 6 parameters for *I* cells and 10 parameters for *E* cells) of each *E*- and *I*-neuron used in intrinsically homogeneous network models. We introduced three different degrees of intrinsic heterogeneities by increasing the extent of jitter.

### Measurements for quantifying patterned activity propagation

As our principal goal here was to assess patterned activity propagation along the *I*- and *E*-rings, we analyzed spike timings from each cell in the network. The instantaneous firing rate for each cell in the network was computed by convolving binarized spike time sequences with a Gaussian kernel with a default standard deviation of 100 ms. These instantaneous firing rates spanning all cells were used to derive metrics that were developed to quantify patterned activity propagation across the rings.

#### Spectral properties of the activity of individual neurons in the network

The first set of metrics measured patterned activity in individual neurons and assessed neuron-to-neuron variability in such patterned activity. The rationale behind these measures was that patterned activity propagation across the network should result in regular firing activity in each neuron in the network. Therefore, when robust activity propagation occurs across the ring, there should be minimal variability across neurons in terms of the power or frequency of activity assessed from different neurons. We computed the power spectrum associated with the instantaneous firing rate of each cell and calculated the maximum power (*P*_4,5_) and the frequency of peak power (*f*_78,9_). The mean, standard deviation, and the coefficient of variation (CV) of *P*_4,5_ and *f*_78,9_ were computed for each cell in the ring were used as measures of robustness of patterned activity propagation. For instance, the CV associated with *P*_4,5_ across cells measured similarity of firing (within a bump of activity) across different cells and the CV associated with *f*_78,9_measured similarity of the frequency at which the propagating activity repeated itself. A lower value for CV associated with *f*_78,9_ indicated that all neurons in the ring were showing patterned firing at the same frequency. A higher value for CV associated with *f*_78,9_reflected the absence of such patterned propagation across cells owing to irregularity in their individual frequency values.

#### Cross-correlation (CC) between firing patterns of different cells in the network

A second set of measurements was derived from the cross-correlograms between a reference cell and all the other cells in the ring. The rationale behind the use of cross-correlograms was to capture the temporal sequence of the progressive propagation of patterned activity along the ring. Specifically, if the ring manifests robust propagation of patterned activity, cells in the ring will fire in the sequence of the organization in the ring. Thus, the temporal shift in the regular firing pattern of cells will be in the higher as they are farther away from each other, with the distance circling back to zero given the ring organization of these cells. In other words, cross-correlations between the activity of a reference cell with all the other cells show a shift in the location where peak cross-correlation (CCpeak) depending on the distance between the two cells under consideration (*e.g.*, Fig. 3*F*). Thus, if the temporal location of the peak of each cell’s cross-correlation (with the reference cell) were plotted as a function of the cell number, stable propagation of patterned activity will manifest as a large positive or a large negative (depending on the direction of activity propagation) slope (*e.g.*, Fig. 3*G*). If, on the other hand, there is no stable propagation of patterned activity across the ring, the slope will be close to zero. Thus, we used the slope associated with the plot of CCpeak *vs.* neuron number as a metric for assessing the manifestation of patterned activity propagation.

Apart from the cell-wise progression associated with the temporal location of CCpeak, we derived metrics associated with the spectra of the cross-correlograms in a manner similar to those we derived from the activity of individual neurons. Specifically, we computed the power spectrum associated with the correlogram of each cell with the reference cell and calculated the maximum power (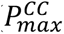) and the frequency of peak power (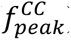). The mean, standard deviation, and the coefficient of variation (CV) of 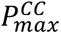 and 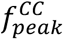 were computed for each cell in the ring and were used as measures of robustness of patterned activity propagation. The CV-based metrics measure the extent of similarity across all cross-correlograms (of the reference cell with each of the other cells). A lower CV would indicate stable propagation of patterned activity across the ring. Threshold values were placed on each of these quantities to identify networks that showed patterned activity propagation across the network (CV of 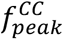 < 0.01 Hz, CV of 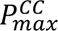 < 0.15, and absolute value of the plot involving slope of shift in CCpeak *vs.* cell number > 0.01).

### Fast HCN channels: Excitability-matched resonance-stripped controls

HCN channels contribute to neuronal dynamics by depolarizing RMP, reducing excitability, and introducing inductive properties to the neuronal membrane through a slow negative feedback loop (Mishra and Narayanan, 2023). The depolarized RMP brings neurons closer to action potential threshold, constituting the mechanism for increasing excitability. On the other hand, the reduction in excitability introduced by HCN channels, through increase in resting conductance and by acting as a negative feedback loop to time-varying signals, reduces both input resistance as well as firing rate. Finally, the introduction of the inductive component manifests as frequency selectivity or intrinsic neuronal resonance in the impedance profile of the neurons. While the depolarizing RMP and the reduced input resistance are not dependent on the slow kinetics of HCN channels, the inductive component (which confers the ability to introduce resonance into neurons) is critically reliant on the slow time course of HCN channels (Hutcheon and Yarom, 2000; Narayanan and Johnston, 2007, 2008). Consequently, an established way of delineating the relative impact of the different biophysical characteristics of HCN channels on cellular and network function is to use a fast version of HCN channels (Narayanan and Johnston, 2007, 2008; Das and Narayanan, 2014; Sinha and Narayanan, 2015). As the voltage-dependent profile of the fast HCN channels remain intact, they still depolarize RMP and reduce input resistance and firing rate, but do not confer resonance upon the neuron that they are expressed in (Hutcheon and Yarom, 2000; Narayanan and Johnston, 2007, 2008; Das and Narayanan, 2014; Sinha and Narayanan, 2015).

To delineate the impact of voltage-dependence *vs*. kinetics of HCN channels on robust propagation of patterned activity in our ring networks, we scaled the time constants of HCN channels in *E* or *I* cells to 5% of their original value. The scaling constructed the fast HCN channels such that they had identical voltage-dependent gating compared to the slow HCN channels with the only difference being in the actual time constants. Expectedly, neurons expressing faster HCN channels lacked membrane potential resonance. As altering the kinetics of HCN channels changes the impedance amplitude of the neurons (Narayanan and Johnston, 2007, 2008; Das and Narayanan, 2014; Sinha and Narayanan, 2015), we tuned the fast HCN channel conductance in each model such that the maximum impedance matched with the value in the neuron model with the slow HCN channels. This additional step ensured that neurons endowed with fast or slow HCN channels were excitability matched.

### Computational details

All single-neuron and network simulations were performed using the NEURON programming environment (Carnevale and Hines, 2006) at 34 °C, with a simulation step size of 25 *μ*s. The data analysis and plotting were executed using custom-written software in the IGOR pro (Wavemetrics). All statistical analyses were carried out using the R (https://www.r-project.org/) computing and statistical package (R core Team 2013).

## ACKNOWLEDGMENTS

The authors thank members of the cellular neurophysiology laboratory for helpful discussions and for comments on a draft of this manuscript.

