## Supplementary Figures for "HCN channels enhance robustness of patterned activity propagation in heterogeneous conductance-based ring networks"

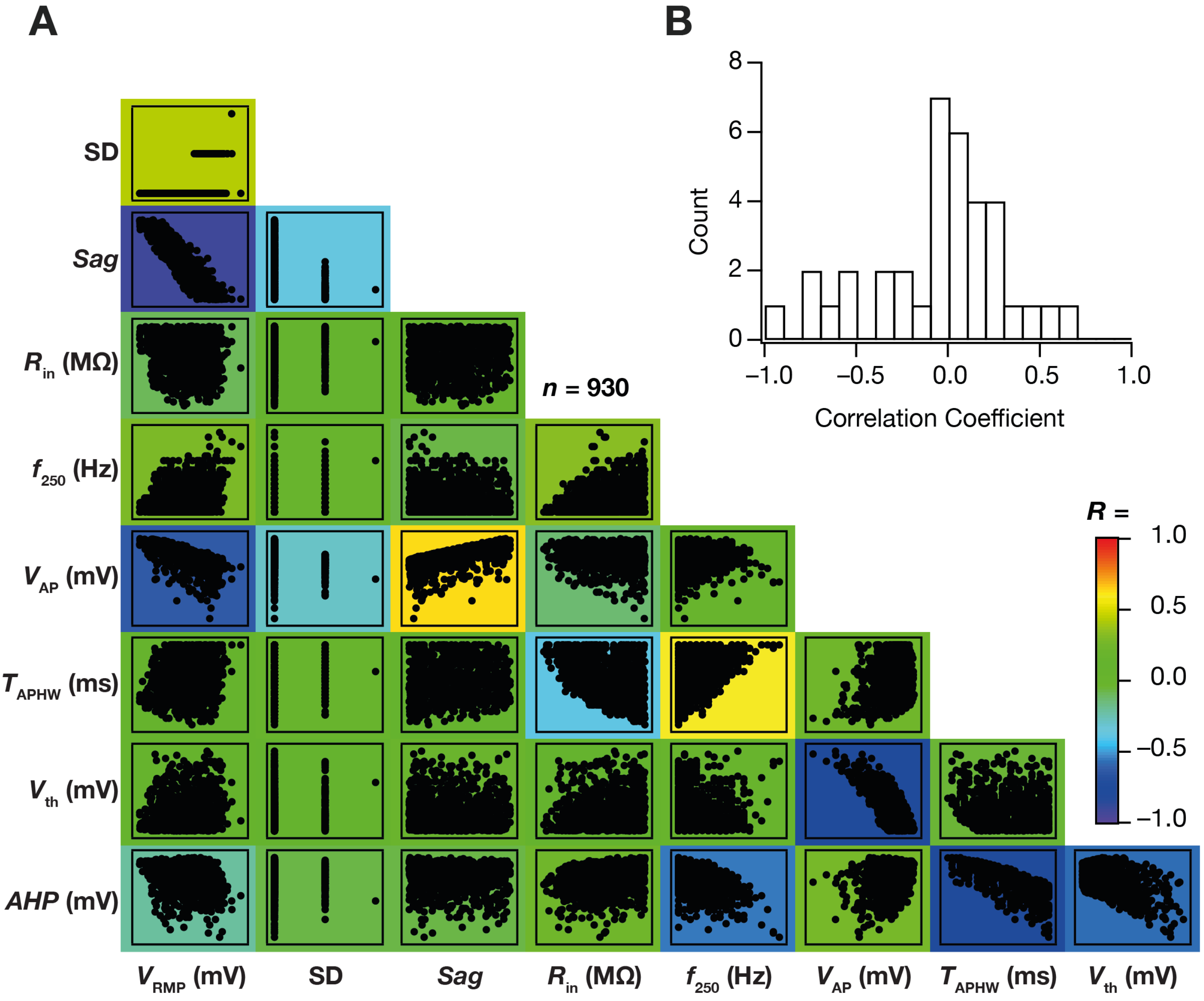

**Figure 1–figure supplement 1: Pairwise correlation across physiological measurements from all valid cortical interneuron models.** (A) Matrix depicting the pair-wise scatter plots (spanning all 930 valid models) between the 9 physiologically relevant measurements, namely $V_{RMP}$, *SD*, Sag ratio, $R_{in}$, $f_{50}$, $f_{250}$, $V_{AP}$, $V_{th}$, $T_{APHW}$, *AHP*. Individual scatter plots are overlaid on a heat map that depicts the pair-wise correlation coefficient computed for that scatter plot. (B) Distribution of the 36 unique correlation coefficient values from scatter plots in A.

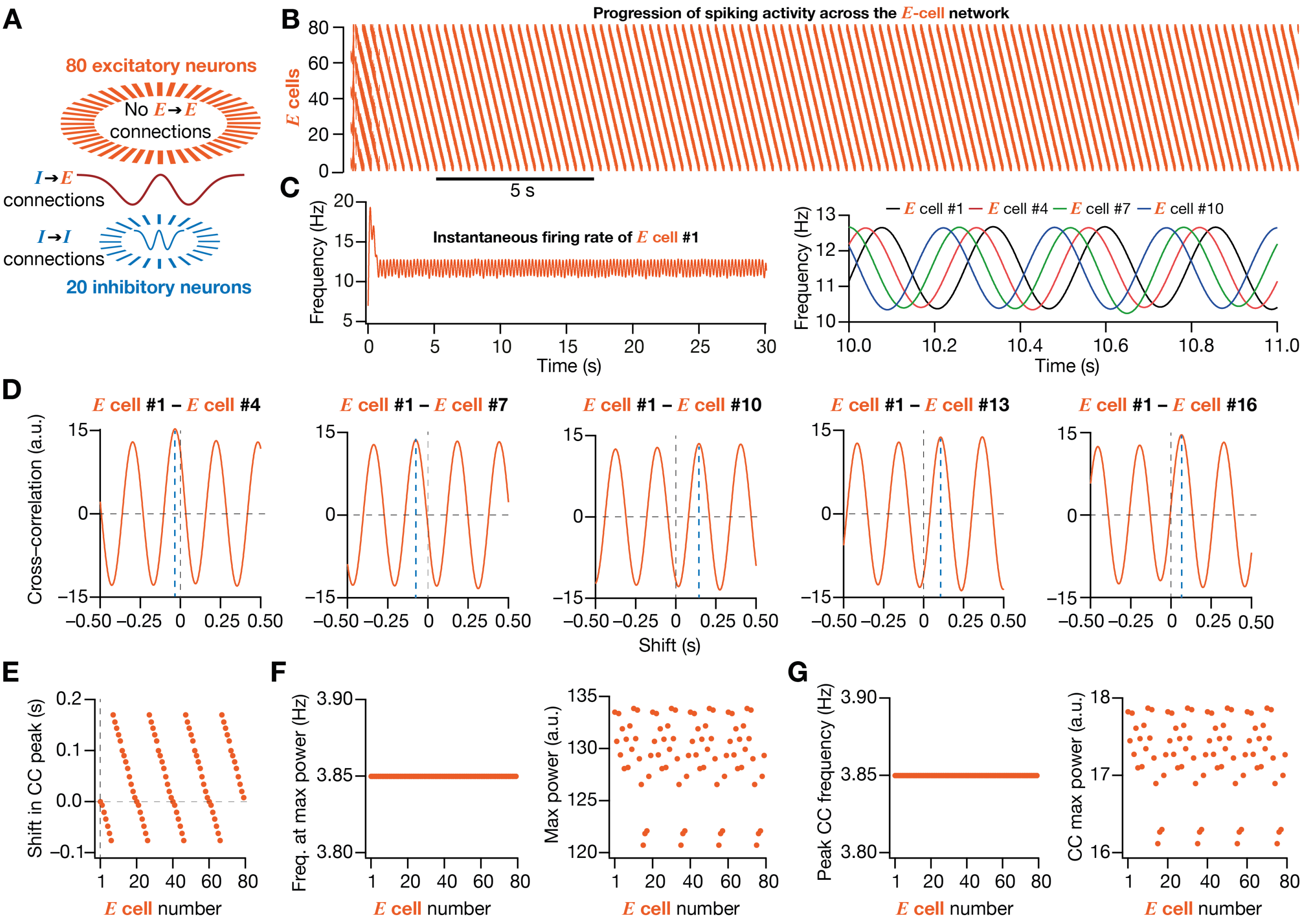

**Figure 3–figure supplement 1: Homogeneous conductance-based ring network model showing patterned activity propagation across *E* cells.** (A) Graphical representation of network architecture with excitatory (*E* cell) and inhibitory (*I* cell) neurons in the ratio of 4:1 (default, 80 excitatory and 20 inhibitory conductance-based model neurons). The synaptic connectivity among inhibitory neurons follows distance-dependent periodic Mexican hat connectivity. Excitatory neurons are not connected with each other. The inhibitory to excitatory neurons connections also follow distance-dependent periodic Mexican hat connectivity. Connections between excitatory neurons to inhibitory neurons were all-to-all. (B) Raster plot showing activity of 80 excitatory neurons exhibiting patterned activity propagation across the ring. Each tick represents a spike in the neuron specified by the column. (C) *Left*, Example instantaneous firing rate of *E* Cell #1, computed by convolving the binarized temporal spike trains (in panel B) of each cell with a Gaussian kernel of standard deviation, $\sigma_{FR}$=100 ms. *Right*, Zoomed version of the instantaneous firing rate of four *E* cells. (D) Example cross-correlogram (CC) between different *E* cells in the network with reference to *E* Cell #1. The progressive shift in the peak of cross-correlogram (shown with blue dotted line) may be noted. (E) Plot showing the time points at which the cross-correlograms (with reference to *E* Cell #1, from panel D) peaked for each *E* cell. This progressive shift in the temporal location of the peak of the cross-correlogram depicts attractor movement over the excitatory neural lattice. (F) Average firing frequency (*Left*) and maximum power (*Right*) from spectral analysis of the activity of each neuron in the network (examples in panel D). (G) Frequency of patterned activity (*Left*), and maximum power (*Right*) computed from spectral analysis of cross-correlograms of *E* Cell #1 with all the other cells (examples in panel D).

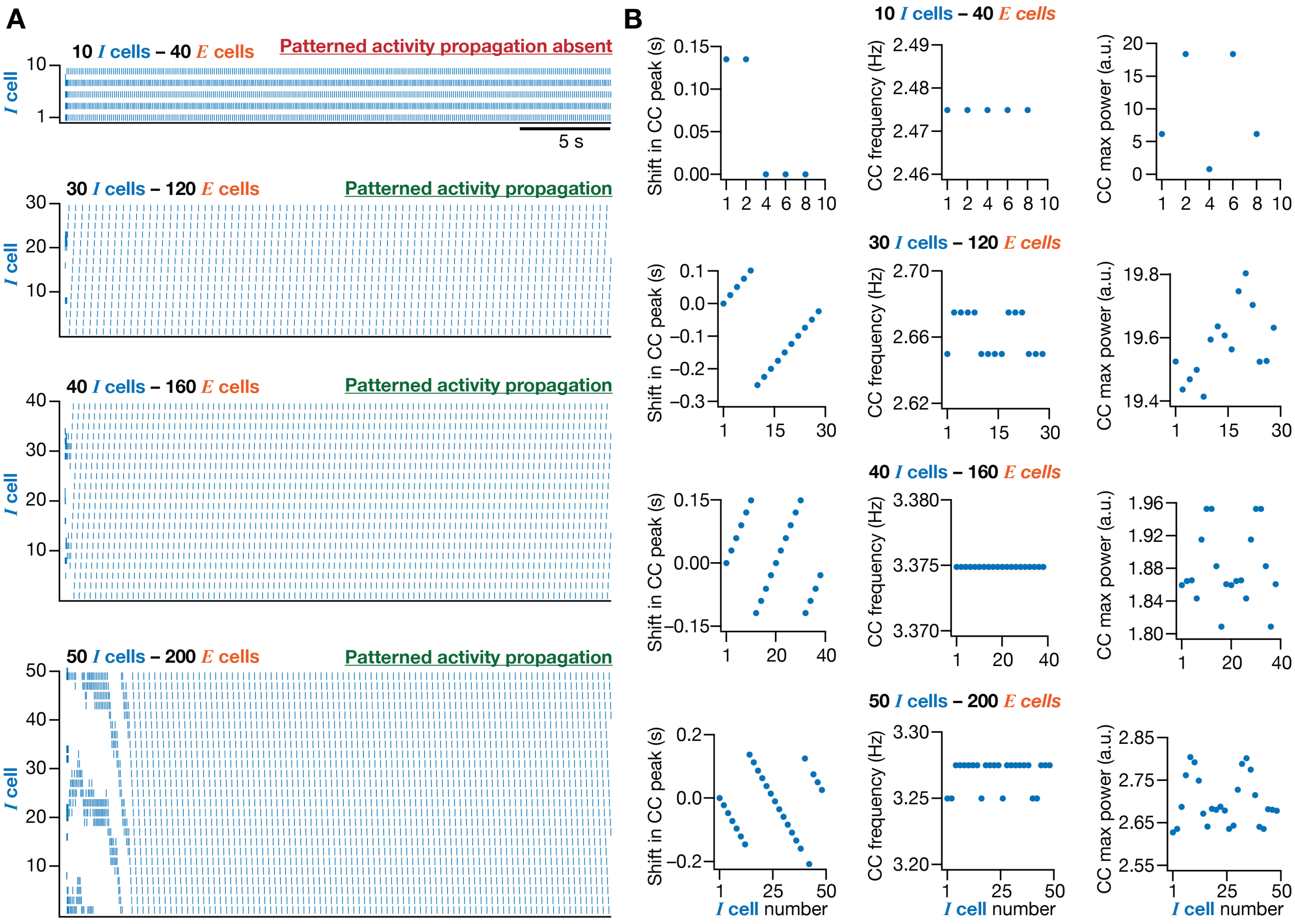

**Figure 3–figure supplement 2: Scalability of patterned activity propagation in homogeneous conductance-based ring network models**. (A) Raster plots for of *I* cells in ring-network models with network size (*E*:*I*), 40:10 (*Row 1*), 120:30 (*Row 2*), 160:40 (*Row 3*), and 200:50 (*Row 4*). Note the absence of patterned activity propagation in the 40:10 network. The other networks showed robust patterned activity propagation. (B) Time points (*Column 1*) at which the cross-correlograms (computed with reference to *I* Cell #1 with all the other cells) peaked as a function of cell number. Firing frequency (*Column 2*) and maximum power (*Column 3*) from spectral analysis of cross-correlograms of *I* Cell #1 with all the other cells for different ring networks shown in panel A.

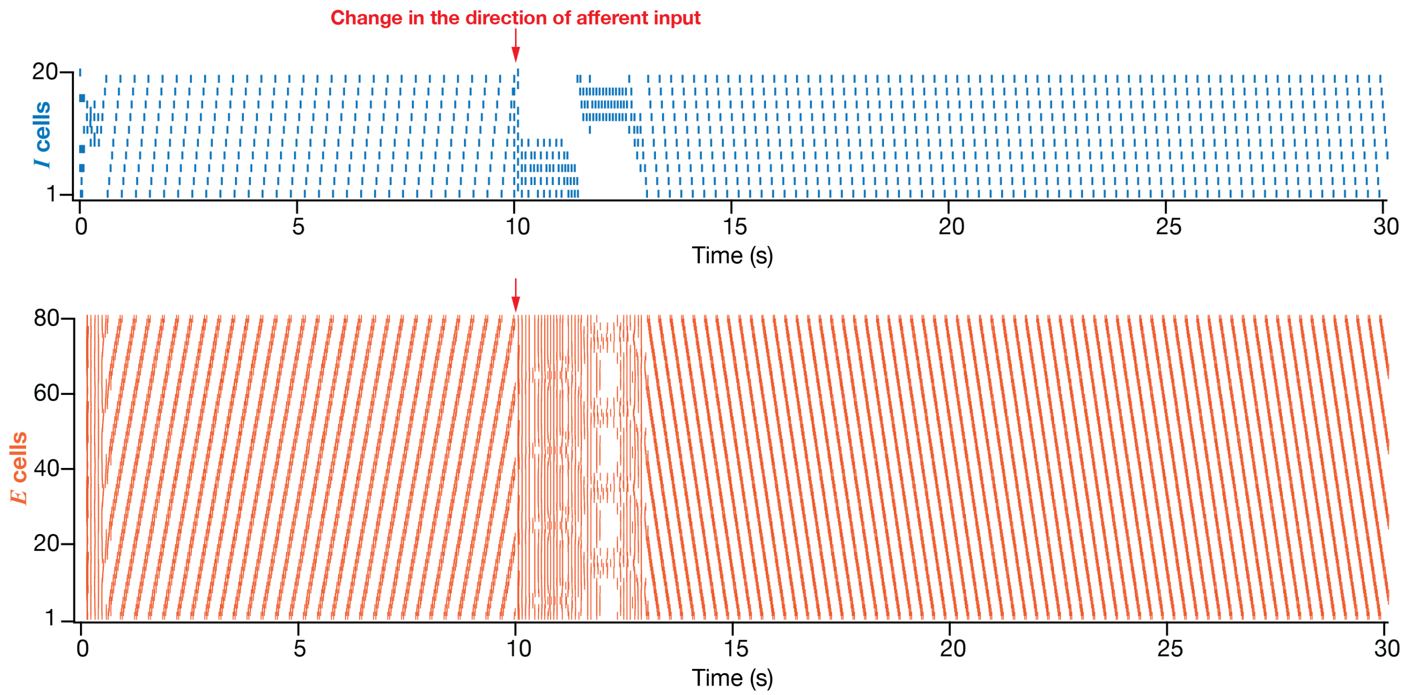

**Figure 3–figure supplement 3: Robust change in direction of patterned activity propagation with change in sign of afferent input.** Raster plots for the activity of inhibitory (*top*; $n = 20$) and excitatory (*bottom*; $n = 80$) neurons manifesting patterned activity propagation across neurons. Note the change in the direction of activity propagation upon a 180° shift in the direction of afferent inputs (introduced at the 10 s time point).

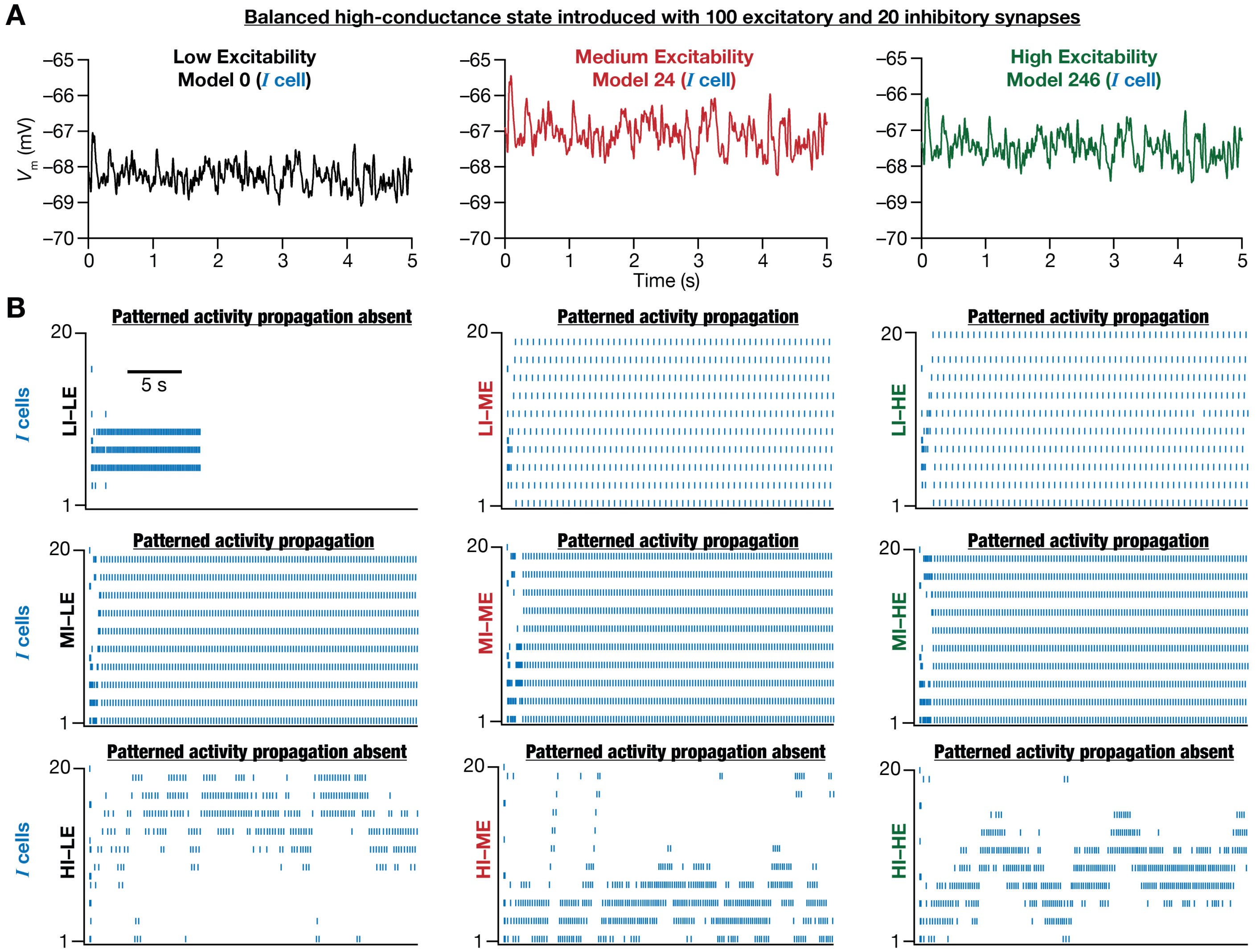

**Figure 4–figure supplement 1: Differential robustness of patterned activity propagation in conductance-based ring network models in the presence of *in-vivo* like high conductance state.** (A) The membrane potential of low (black), medium (red), and high (green) excitability inhibitory neurons with *in vivo* like high conductance state (HCS). (B) Raster plots for the 9 network models used in Figure 4 with HCS introduced in each neuron of the network. Note for some networks with specific combinations of inhibitory and excitatory neurons, patterned activity propagation was completely lost. However, patterned activity propagation was achievable in these models with retuning of network parameters.

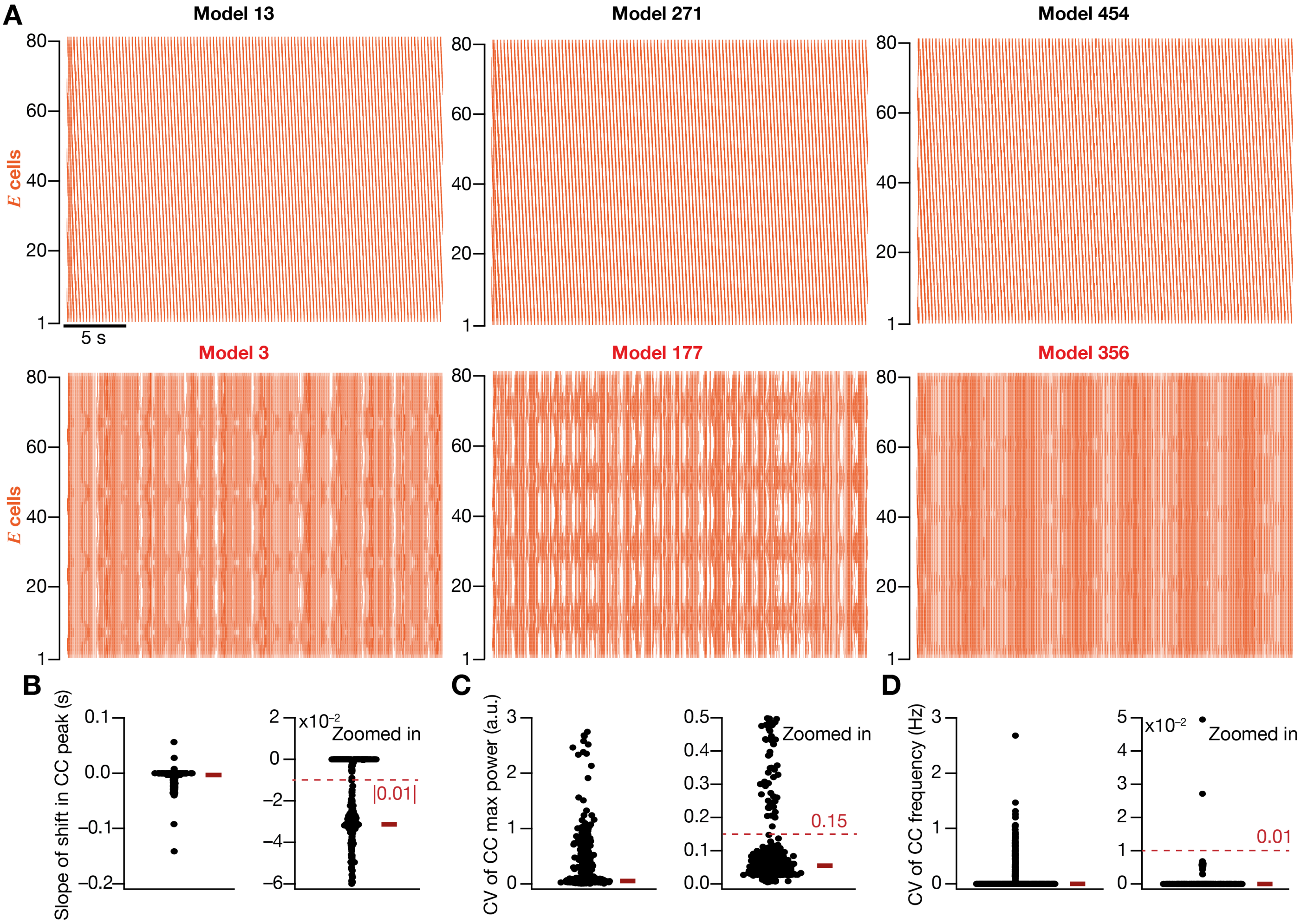

**Figure 5–figure supplement 1: Distribution of measurements from randomized homogeneous ring networks.** (A) *Row 1,* Raster plots of example network models exhibiting robust propagation of patterned activity. *Row 2,* Raster plots of example network models where propagation of patterned activity was absent. Shown are the activity patterns of E-cells ($n$ = 80). (B) *Left*, Linear slope of the shift in peak cross-correlogram (CC) between "E Cell #1" and all other cells in the excitatory layer ($n$ = 500 randomized networks: 5 trials each on 100 networks). *Right*, Zoomed-in version of the plot on the left, also showing the threshold for declaring valid models based on patterned activity propagation across the E-cell network. (C–D) Same organization as (B), but for coefficient of variance (CV) for maximum power (C) and peak frequency (D) computed across all E cells. While a lower threshold on the slope of shift in the CC peak provided a measure of propagation of activity across the network, upper thresholds on the variability in maximum power and peak frequency ensured that the dynamics was stable across all E cells. Networks that manifested a minimum slope of shift (B) and were within the limits of maximally allowed variability in power (C) and frequency (D) were declared as valid models.

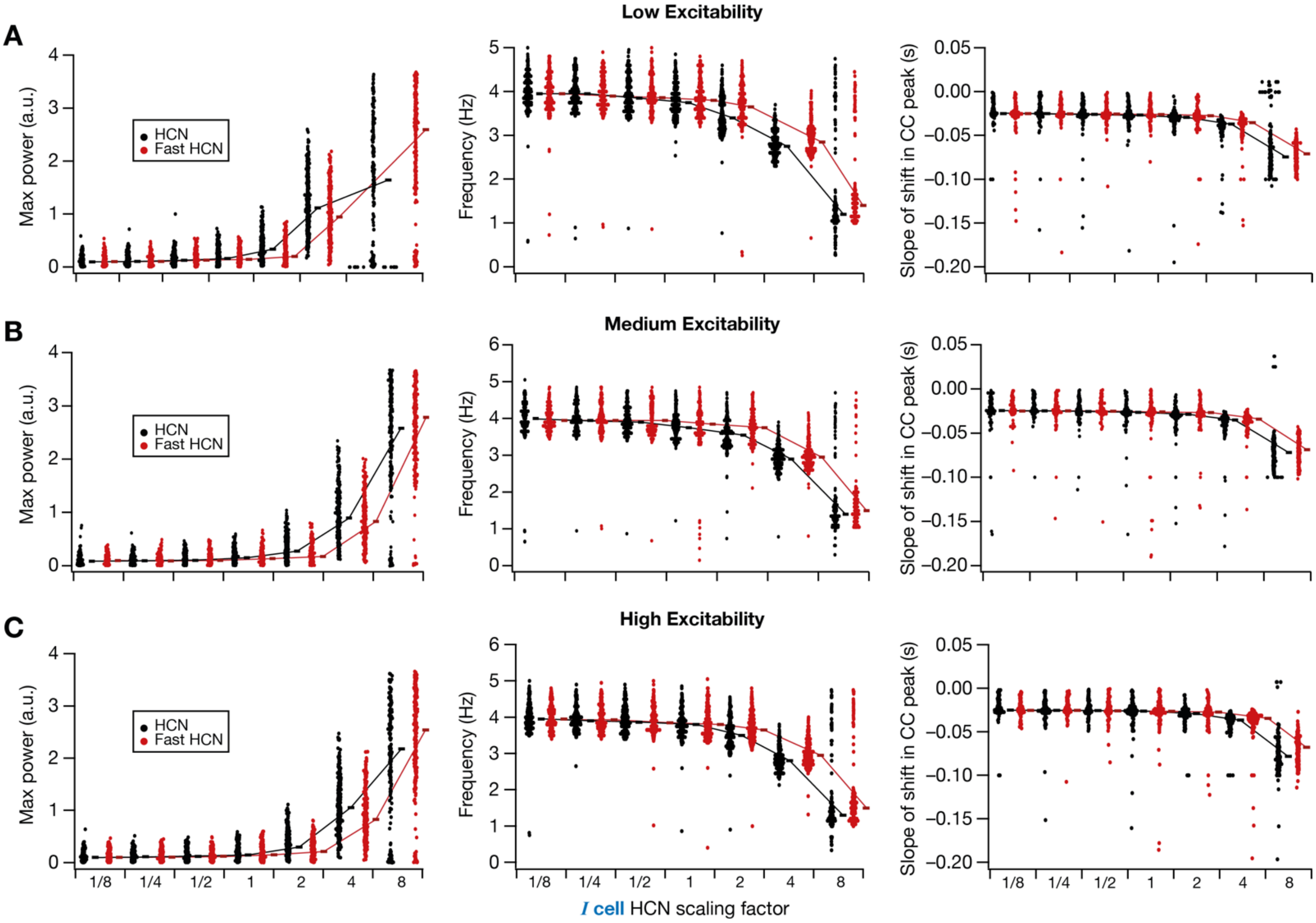

**Figure 6–figure supplement 1: Impact of HCN channels modulation on measurements across all valid homogeneous ring networks.** (A) Mean maximum power (*Left*), mean firing frequency (*Center*), and linear slope of the shift in cross-correlogram peak (*Right*) for each valid network constructed with low excitability *I* cells. Shown are values for different fold-changes in networks endowed with HCN or Fast HCN channels. (B–C) Same as (A), for ring networks obtained with medium (B) or high (C) excitability *I* cells. The black/red rectangle adjacent to each plot represents the respective median values, and are connected black/red lines to illustrate the dependencies.

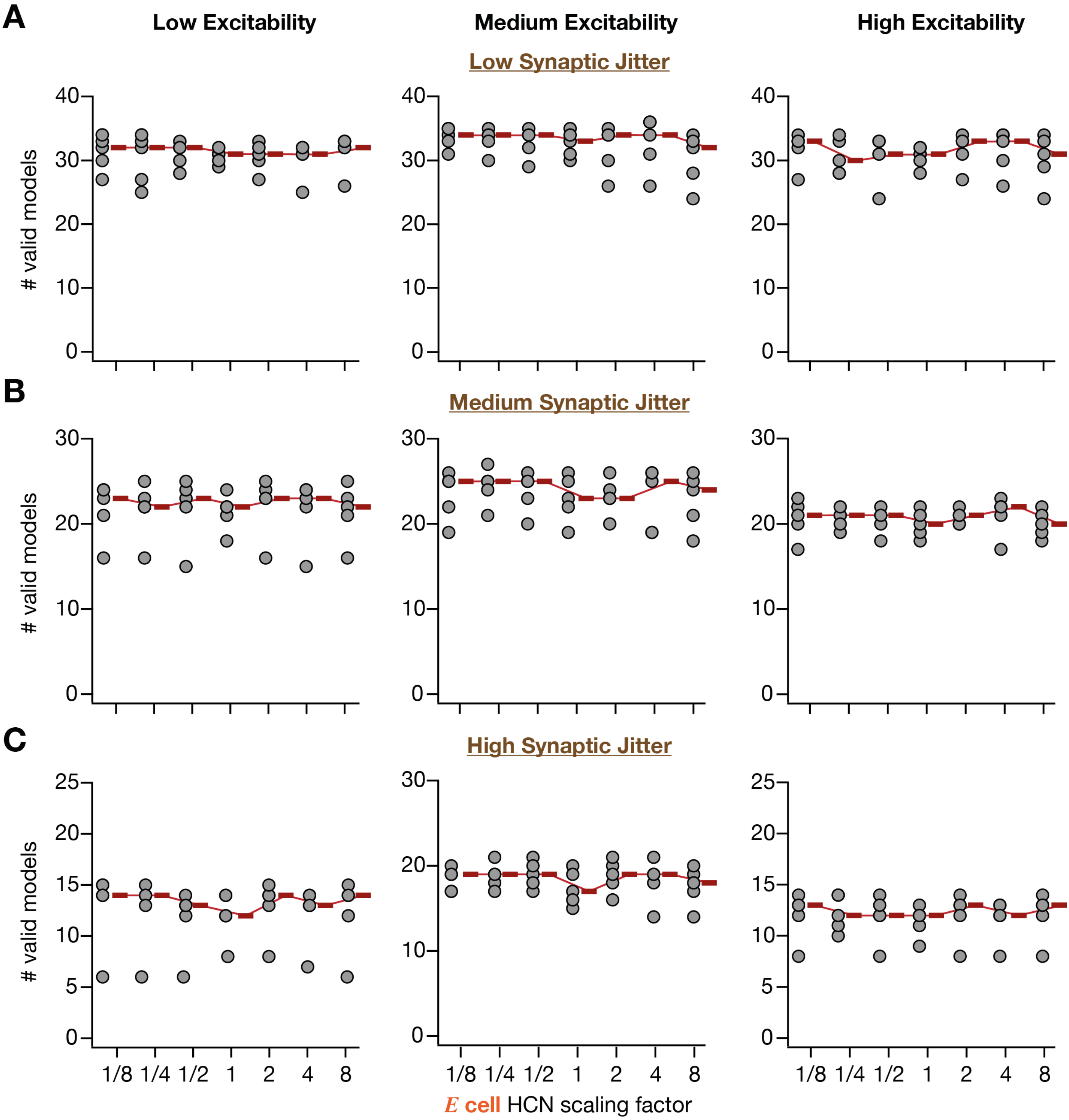

**Figure 7–figure supplement 1: Modulation of HCN channel of excitatory neurons did not stabilize patterned activity propagation in synaptically heterogeneous ring networks.** (A) Number of valid network models (out of 50 models for each trial, across 5 independent trials) plotted as a function of HCN conductance of the *E* cells for networks with all three levels of excitability. These networks were perturbed with a low degree of synaptic jitter. (B–C) Same as (A) for medium (B) and high (C) degree of synaptic jitter. The red rectangle adjacent to each plot represents the respective median value.

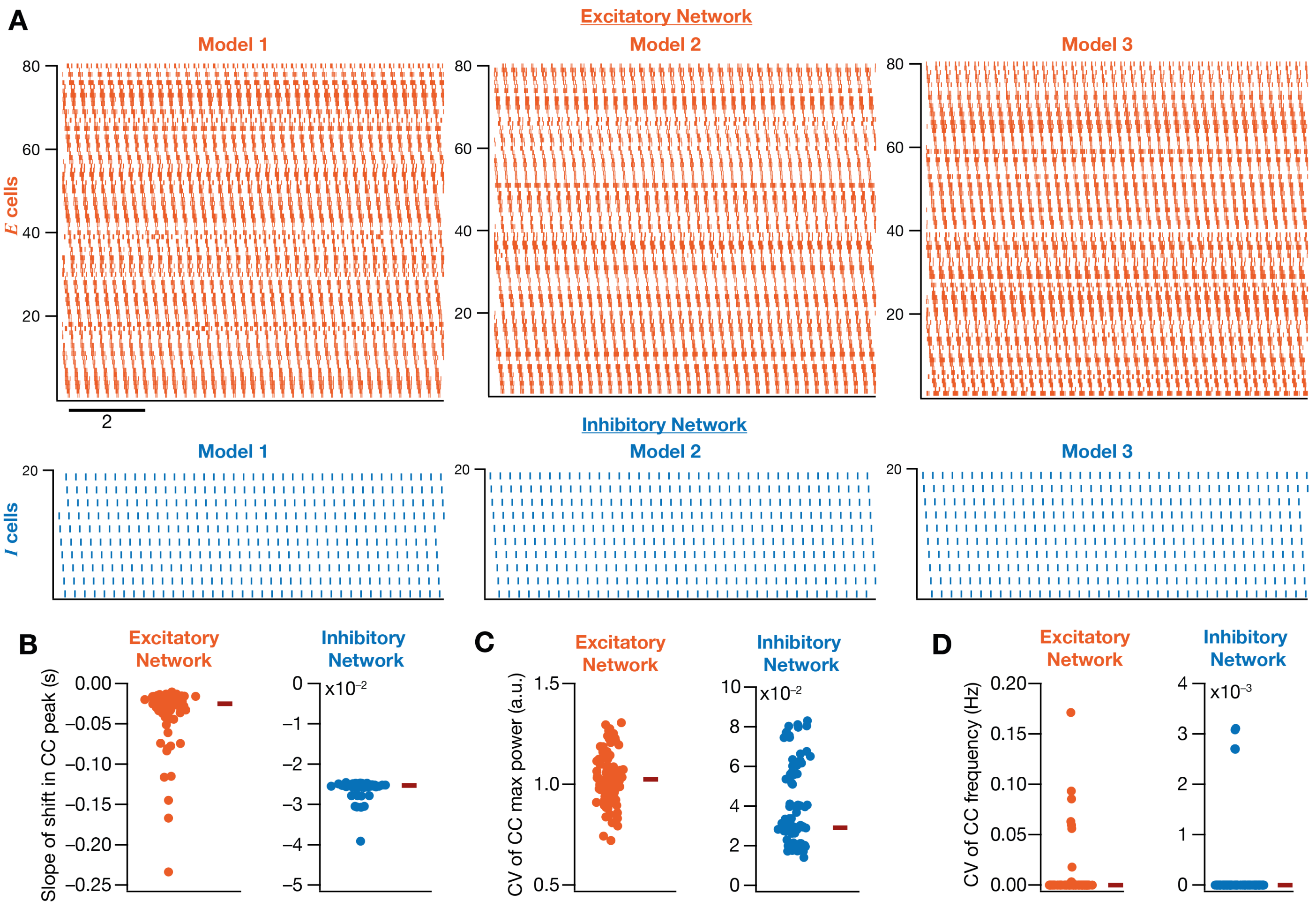

**Figure 8–figure supplement 1: Ring networks with intrinsic heterogeneities confined to E cells showed robust propagation of patterned neural activity.** (A) Examples of raster plots of networks manifesting robust propagation of patterned activity in their excitatory (*Top*) and inhibitory (*Bottom*) layers. Each *E* cell in each of these networks was unique, thus making the E-cell network heterogeneous. The inhibitory layer was homogeneous (same *I* cell repeated) and was constructed from a medium excitability *I* cell. (B) Linear slope of the shift in peak cross-correlogram (CC) between E Cell #1 and all other cells in the excitatory (*Left*; $n$ = 100 models) and inhibitory (*Right*; $n$ = 100 models) neurons. (C–D) Coefficient of variance (CV) for maximum power (C) and peak frequency (D) computed for excitatory and inhibitory cell layers. The red rectangles adjacent to each plot in (B–D) represent the respective median values.

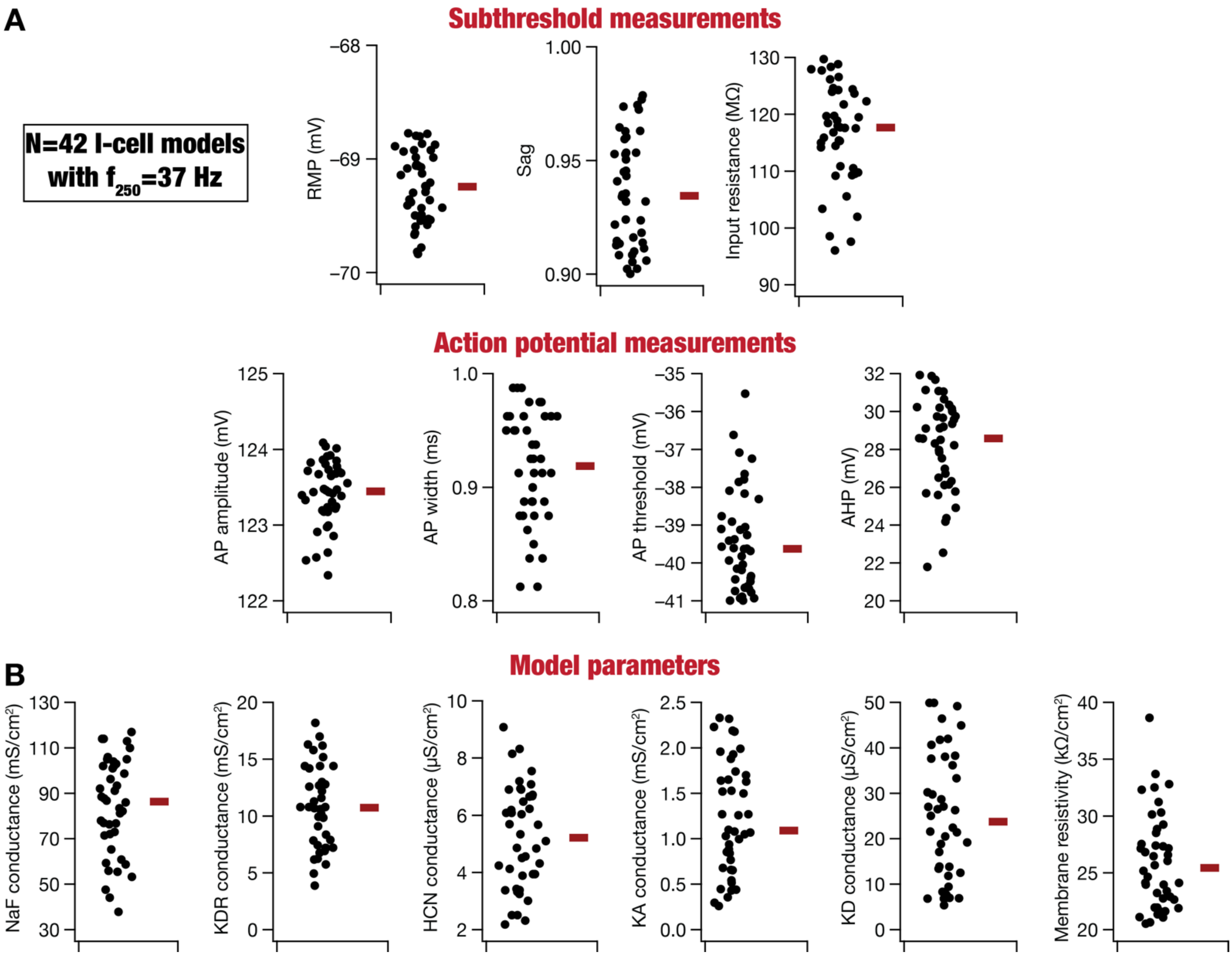

**Figure 8–figure supplement 2: Heterogeneities in intrinsic measurements and model parameters of all *I* cells with $f_{250}$ at 37 Hz.** Measurements (A) and parameters (B) of all *I*-cell models with their $f_{250}$ at 37 Hz (medium excitability *I* cells) manifested pronounced heterogeneity (*cf.* Tables 1–2 for ranges of parameters and measurements). *N*=42 for all panels.

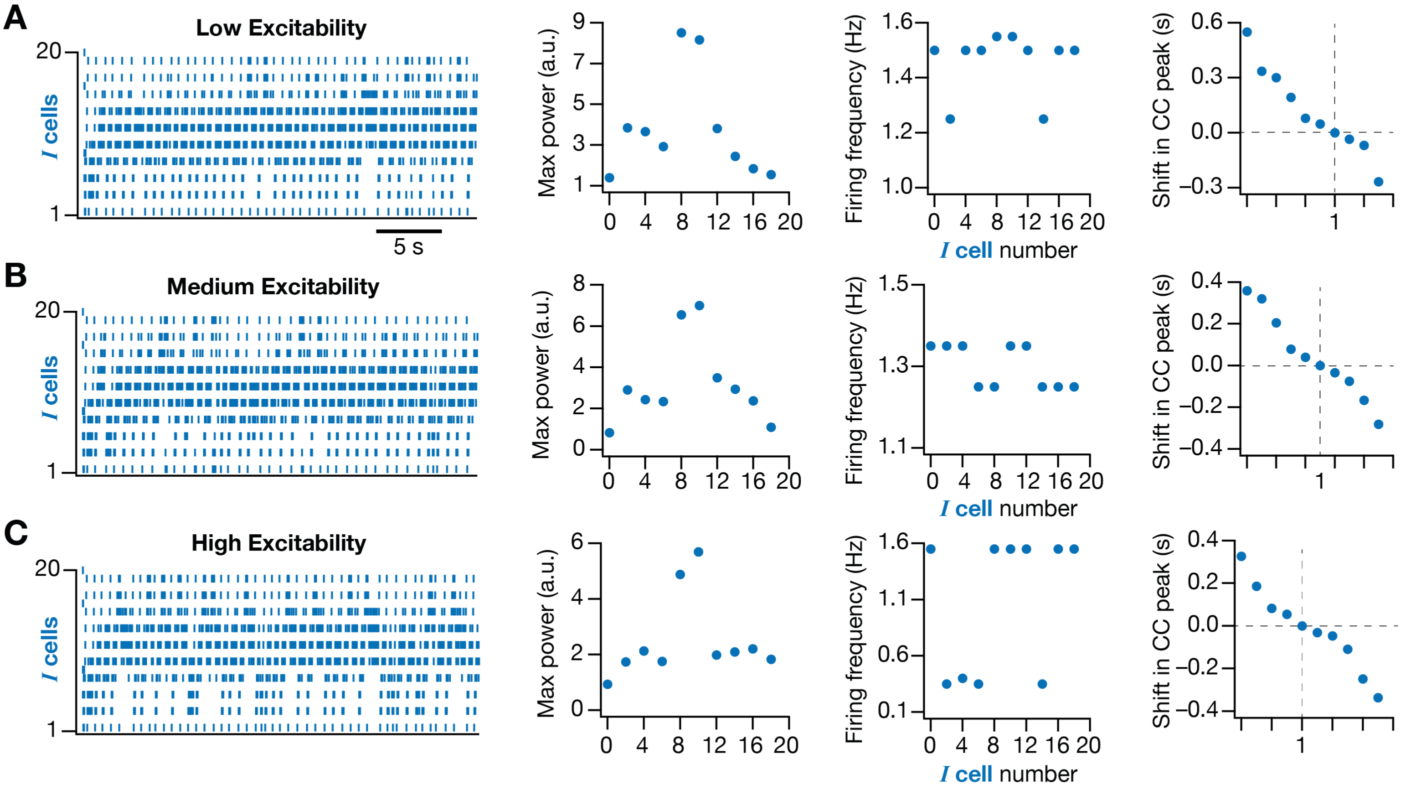

**Figure 8–figure supplement 3. Ring network models with limited intrinsic heterogeneity in inhibitory neurons showed robust propagation of patterned neural activity.** (A) Example raster plot (*Row 1*), average maximum power (*Row 2*), average firing frequency (*Row 3*), and shift in the cross-correlogram (CC) peak (*Row 4*) plotted for each I cell in a ring network. The ring network was constructed with *I* cells showing limited heterogeneity: $f_{250}$ for all the inhibitory neurons in the network was 37 Hz, but all the other intrinsic properties were heterogeneous across *I* cells. This network contained low-excitability *E* cells. (B–C) Same as (A) but with medium and high excitability for *E* cells.

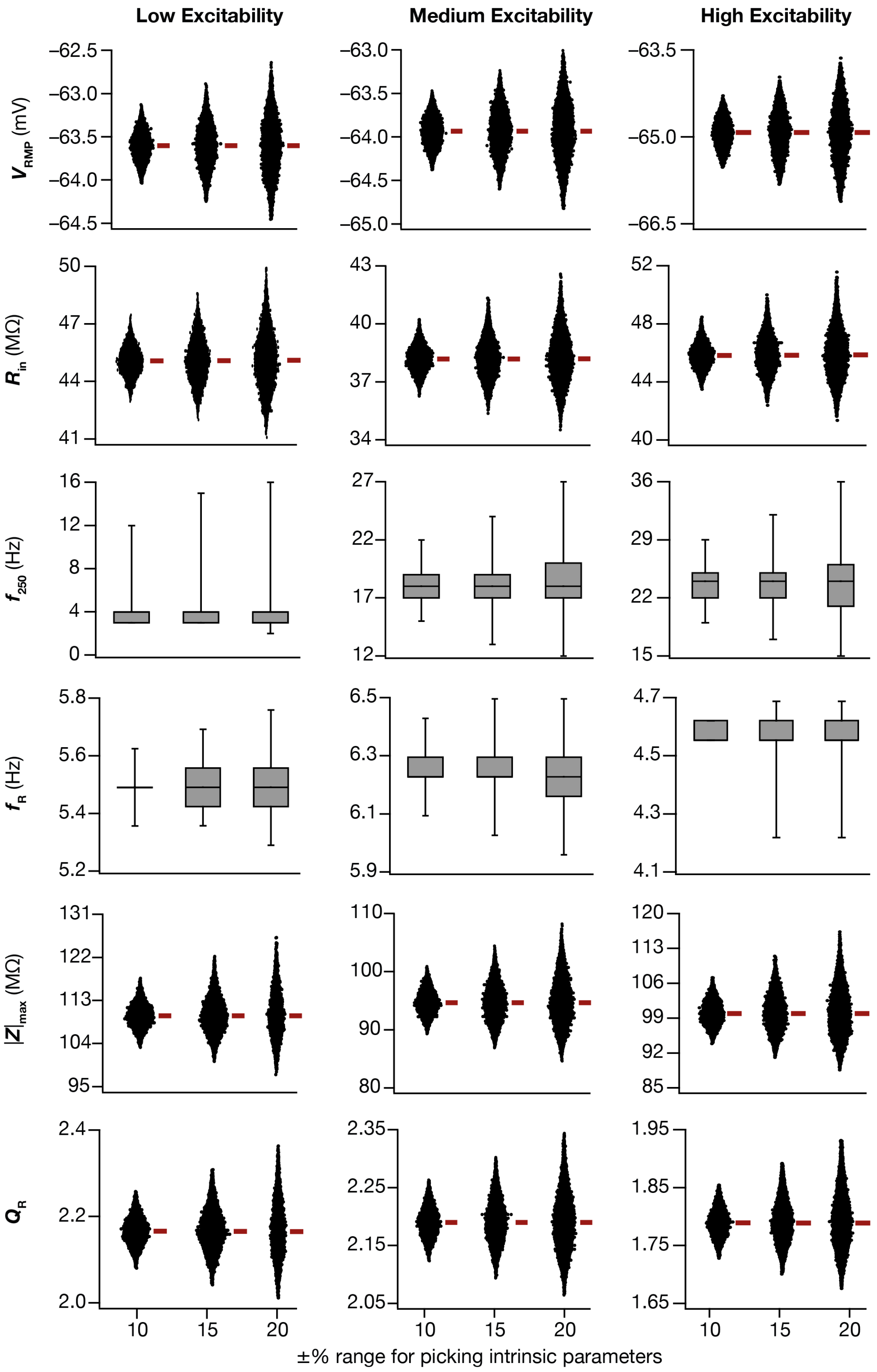

**Figure 8–figure supplement 4: Intrinsic properties of excitatory neurons with different degrees of intrinsic heterogeneities.** (A) Values of 6 intrinsic measurements (resting membrane potential ($V_{RMP}$), firing frequency at 250 pA current injection ($f_{250}$), input resistance ($R_{in}$), resonance frequency ($f_R$), impedance at resonance frequency ($|Z|_{max}$), and strength of resonance ($Q$)) for different degrees of intrinsic heterogeneities in excitatory neurons with low (*Left*), medium (*Center*), and high (*Right*) excitability neurons. The red rectangle adjacent to each plot represents the respective median value. $n$ = 20000 (250 network models with 80 $E$ cells each) for each degree of intrinsic heterogeneity.

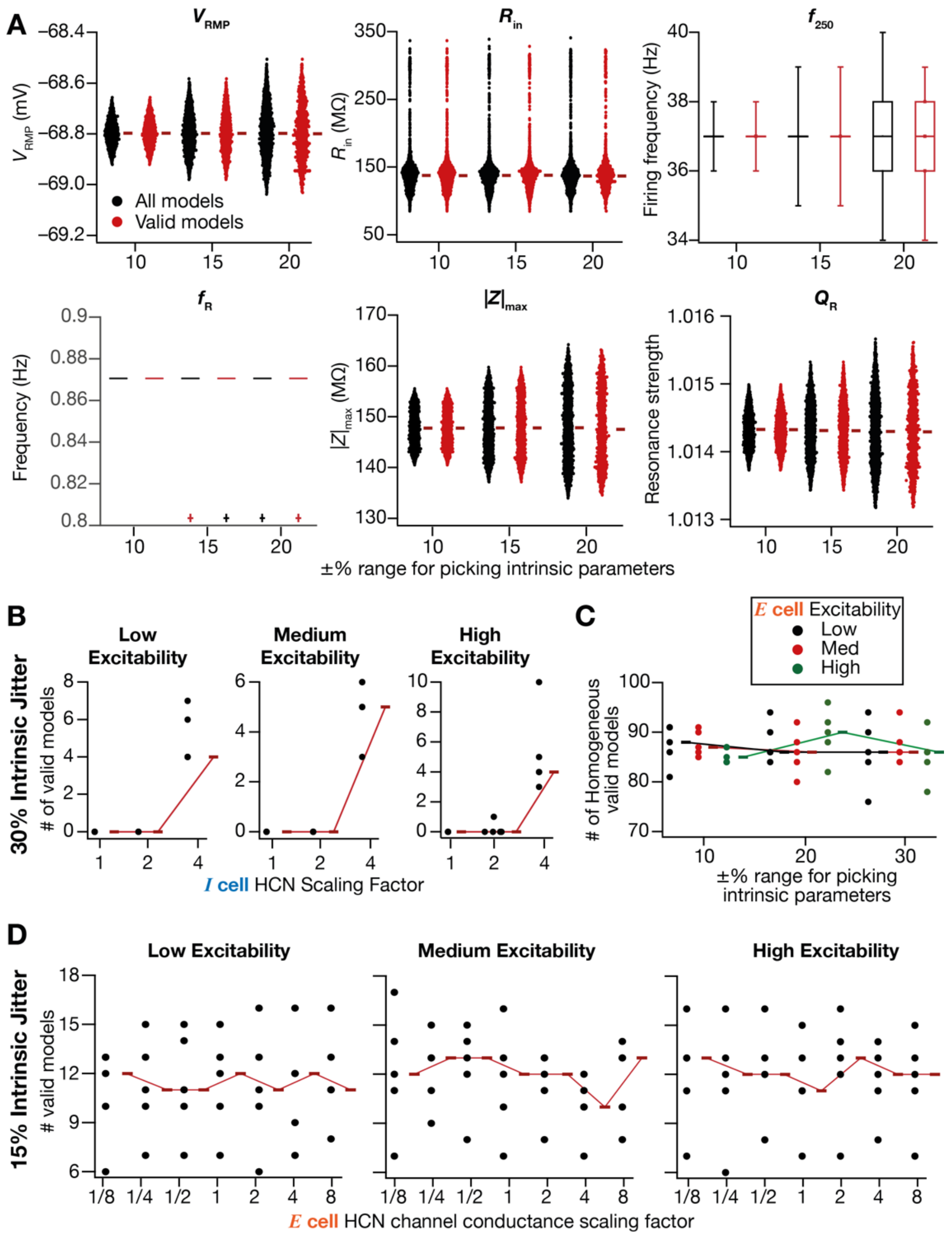

**Figure 8–figure supplement 5. Intrinsic heterogeneities are stabilized by HCN channels in CAN models.** (A) Values of intrinsic measurements (resting membrane potential ($V_{RMP}$), firing frequency at 250 pA current injection ($f_{250}$), input resistance ($R_{in}$), resonance frequency ($f_R$; although outliers are shown separately, resonance was absent in all these cases with $f_R < 1$ Hz), impedance at resonance frequency ($|Z|_{max}$) and strength of resonance ($Q$)) for different degrees of intrinsic heterogeneities for inhibitory neurons of all models (black, 250 network models with 20 neurons each, $n = 5000$) and valid models (red, 10% intrinsic jitter ($n = 247 \times 20 = 4940$), 15% intrinsic jitter ($n = 214 \times 20 = 4280$), 20% intrinsic jitter ($n = 117 \times 20 = 2340$)) obtained with four-fold change in HCN conductance in low excitability case. The distributions associated with valid models cover the entire range suggesting a lack of restriction on intrinsic properties to obtain robust propagation of patterned activity. (B) Number of valid models plotted a function of fold-change in HCN channel conductance in inhibitory neurons of the network with low (*Left*), medium (*Center*), and high (*Right*) excitability neurons for neurons endowed with 30% intrinsic jitter. A four-fold increase in HCN channels conductance in inhibitory neurons was able to make 4–10 ring networks (out of 50 models) exhibit robust propagation of patterned activity. (C) Robustness of the patterned activity propagation in homogeneous ring networks. The intrinsic properties of the neuron were chosen from 10%, 20%, or 30% intrinsic jitter for all three levels of excitability. (D) Number of valid models plotted a function of fold-change in slow HCN channel conductance in *E* cells of the network with low- (*Left*), medium- (*Center*), and high-excitatbility (*Right*) *E* cells with 15% intrinsic jitter. The red rectangle adjacent to each plot represents the respective median value.
